# In-depth phenotyping reveals unexpected floral trait variation in *Mimulus cardinalis* across a range-wide latitudinal gradient

**DOI:** 10.1101/2025.10.03.680188

**Authors:** Mikhaela Neequaye, Elenor B. Kennedy, Hannah Gunn, Katherine E. Wenzell, Kelsey J.R.P. Byers

## Abstract

**Background & Aims:** Pollinators play a crucial role in the evolution and diversification of flowering plants. Intraspecific variation in floral traits occurs frequently and likely reflects variable biotic and abiotic selection pressures. The hummingbird-pollinated monkeyflower *Mimulus cardinalis* occurs across western North America, where geographic mosaics of selection have likely influenced its natural history, due to this wide species range. Processes driving early stages of divergence and speciation remain poorly understood, though theory predicts that trait divergence will be most likely at range edges. Within *M. cardinalis,* which is usually red-flowered and hummingbird-pollinated, two independent shifts to yellow flowers have occurred at the northern and southern range edges, including a population found on Cedros Island, off the coast of Baja California, Mexico.

**Methods:** This study characterises the local climatic variables of five accessions of *M. cardinalis* derived from locations across a latitudinal geographic gradient. Highly integrative methods in metabolomics and morphological analyses were used to characterise a suite of pollinator-relevant floral traits across the range of *M. cardinalis* species variation. We use transcriptomics and whole genome sequencing to study underlying genetic variation in two yellow-coloured, range-edge accessions.

**Key Results:** This study uncovers high levels of variation in morphology, nectar properties, pigmentation and scent profile across geographically diverse *Mimulus cardinalis* accessions, in addition to profiling how these accessions are perceived by potential pollinators. We find high levels of phenotypic variation across *M. cardinalis*, particularly in the biochemistry of pigment and scent, where inflorescences that appear to be the same shade of red have completely different anthocyanin profiles. Floral trait-underlying genetic differences between different *M. cardinalis* lines were also investigated, revealing potential mechanisms underlying floral diversification.

**Conclusions:** This work highlights the importance of interplay between floral trait diversification, pollinator perception and climate. This work also highlights the importance of considering suites of floral traits in a quantitative fashion when studying intraspecific variation in relation to species range. This study also provides genetic insight into changes in these traits and provides many future directions to study the early stages of pollinator-mediated trait differentiation across a species range.

## INTRODUCTION

Shifts between different animal pollinators often coincide with speciation events in flowering plants (Forest et al., 2013). Different pollinators often prefer different combinations of floral traits, which can lead to preferential visitation of a specific subset of the population with more favourable traits, promoting reproductive isolation and subsequent speciation (Kay and Sargent, 2009). Floral traits are important for reproduction and species cohesion, but intraspecific variation in floral traits, both continuous and discrete, is common (Herrera et al., 2006; Westerband et al., 2021; Petren et al., 2021; Wenzell et al., 2021; Wenzell et al., 2023). Such variation likely reflects responses to different selection pressures and plasticity under varying environmental and ecological conditions (Kuppler et al., 2020). These variable conditions can generate geographic mosaics of selection, which can drive local adaptation and potentially allopatric speciation when divergent selection acts on geographically separated populations (Thompson et al., 2005)

Processes driving early and ongoing stages of divergence and speciation remain poorly understood, though theory predicts that trait divergence will be most likely at range edges due to decreased gene flow and thus increased isolation as well as distinct ecological conditions relative to the centre of the range (Kay and Sargent, 2009). This becomes particularly acute when considering recent angiosperm introductions to islands, which have their own biomes, ecology and both reproductive and evolutionary pressures (Bremer & Eriksson, 1992) and are often limited in potential pollinating fauna compared to mainland areas, leading to potential pollen limitation and a switch to generalist pollination (Zell et al. 2024).

Because selection mediated by pollinators plays a central role in angiosperm evolution and diversification, pollinator interactions have received much attention as drivers of floral trait variation (van der Neit and Johnson 2012). Nonetheless, floral traits are increasingly recognized to reflect selection and plastic responses to abiotic conditions as well (Strauss & Whittall 2006, Coberly and Rausher, 2003). Abiotic factors such as water availability are known to influence floral pigment (Gardiner & Glennon, 2019) as well as floral size (Caruso et al., 2019; Galen, 2005) and scent (Cna’ani et al., 2014; Luizzi et al., 2021). Examining multiple potential drivers of floral trait evolution in a geographic context is needed to better understand how divergence may arise at early stages, particularly as biotic (e.g. pollinators) and abiotic drivers are likely to act in combination with each other (Johnson et al., 2006).

*Mimulus* sect. *Erythranthe* is an emerging model for pollinator-mediated floral evolution (Yuan, 2019). Within section *Erythranthe*, there are thought to have been two independent shifts from insect-mediated pollination to hummingbird-mediated pollination (Beardsley et al., 2003), although significant phylogenetic discordance exists (Nelson et al. 2021). This correlates with a transition to hummingbird-related floral traits, such as red pigmentation, more reflexed petal lobes, tubular corollas and more abundant nectar (Beardsley et al., 2003, Wenzell & Neequaye et al., 2025).

Floral scent is understudied relative to other floral traits (Byers, 2021), but it has been shown that floral volatile organic compounds (VOCs) also contribute to pollinator preference and pollinator-mediated speciation. For example, key scent compounds contribute to the attraction of bees to bee-pollinated *Mimulus lewisii*, compared to its hummingbird-pollinated sister species *M. cardinalis* (Byers et al., 2014a). Meanwhile, hummingbird-pollinated flowers typically have lower scent emission than insect-pollinated flowers, linked to hummingbirds’ poor scent perception and limited scent memory (Byers et al., 2014b, Knudsen et al. 2004, Goldsmith & Goldsmith 1982, Núñez et al., 2021).

*Mimulus cardinalis* is a perennial herb found to occur in a latitudinal gradient across western North America, from central Oregon (USA) to northern Baja California (Mexico) (Angert et al., 2018). It is unlikely that *M. cardinalis* experiences uniform biotic or abiotic selection pressures across this wide geographic range, where interannual variability in climatic factors such as temperature and precipitation is particularly acute in the southern range (Wooliver et al., 2020). While *M.cardinalis* is typically hummingbird-pollinated, with the aforementioned hummingbird-associated traits (Fig. 1), stable yellow-flowered populations have arisen independently at the northern and southern *M. cardinalis* range edges. This may influence pollinator preference, as insect pollinators such as bumblebees and hawkmoths prefer yellow variants to red variants in *Mimulus* section *Erythranthe* (Vickery and Vickery, 1992, Wenzell & Neequaye et al., 2025, Byers & Bradshaw 2021). There is evidence supporting assortative mating within the yellow-flowered morph, mediated by pollinator fidelity (Vickery, 1995), suggesting that these changes could lead to reproductive isolation and speciation between red- and yellow-flowered populations.

**Figure 1.**
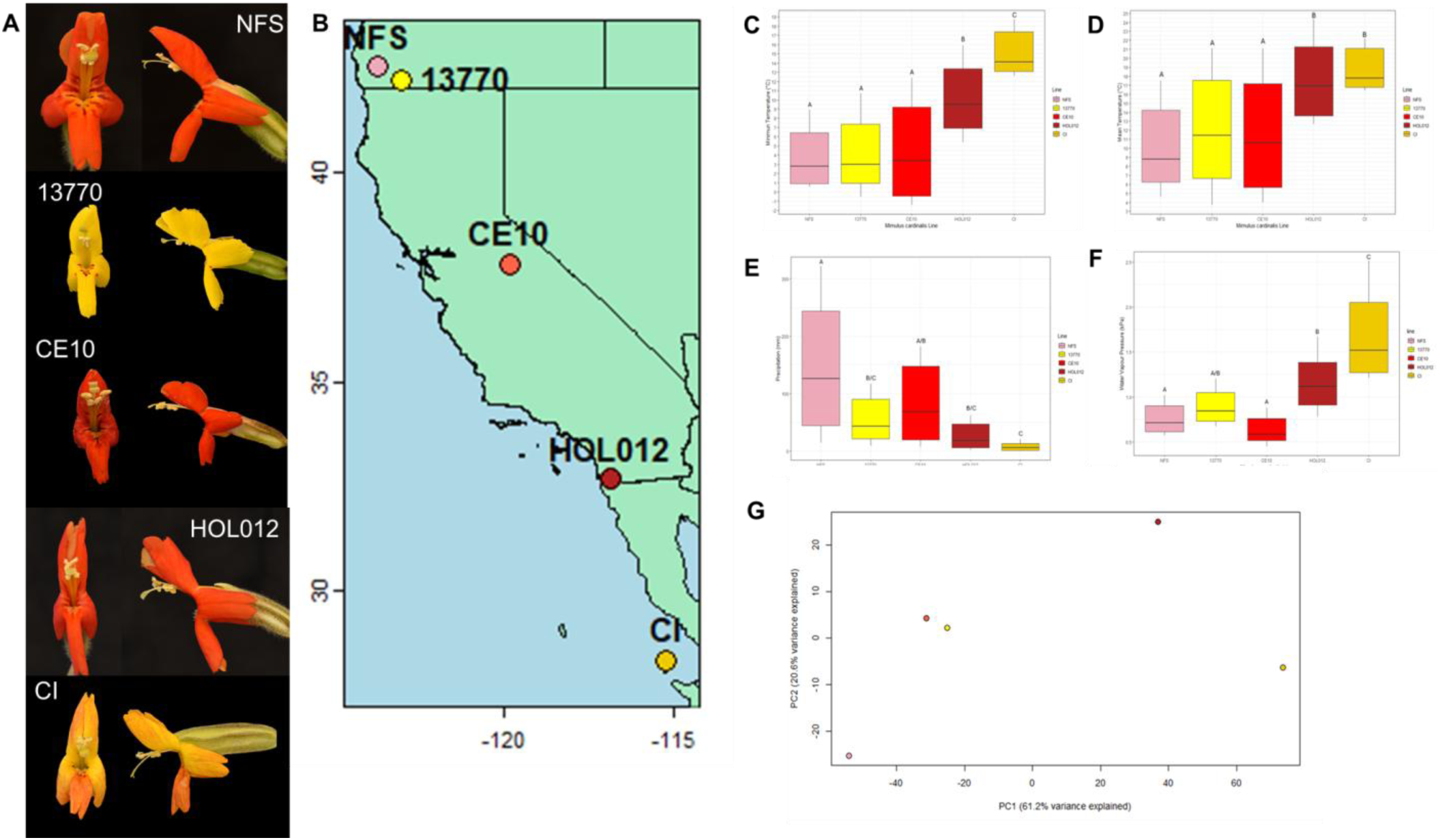
Focal morphs and climate. (A) Front and side images of *M. cardinalis* accessions derived from a latitudinal gradient across Western North America. (B) Map of seed origins for accessions. (C-G) Climate variables: (C) Minimum temperature, (D) Mean Temperature, (E) Precipitation, (F) Water vapour pressure and (G) PCA of all climate variables. Climate data is based on best estimates of seed sampling locations (B). Variance shows a 30-year average from 1970-2000. Significance values were obtained using one-way ANOVA, followed by Tukey’s HSD for pairwise comparison. Bars sharing a letter are not significantly different, and lettering is independent for each variable. Bars show median expression, while error bars represent mean ± standard deviation.

One of these yellow-flowered *M. cardinalis* populations is found at the northern range edge, in the Siskiyou mountains, Oregon, USA. This population was characterised in (Wenzell & Neequaye et al., 2025) and is referred to here as accession 13770. Wenzell & Neequaye et al., 2025 found flowers of 13770 to have a shorter corolla with a wider opening, increased carotenoid content, reduced anthocyanin content, increased total scent emission and a different scent profile to those of the red-coloured, range-centred CE10 accession. These large and small-scale transitions in floral traits contribute to increased attraction of insect pollinators, like bumblebees and, potentially, hawkmoths (Wenzell & Neequaye et al., 2025). The other population of yellow-flowered *M. cardinalis* is found at the southern range edge, on Cedros Island, Baja California, Mexico, is referred to here as accession CI. Cedros Island represents the most biodiverse of the islands in the Baja California Pacific, containing 266 plants of which 14 are endemic (Ratay et al., 2014). It is also believed to have been connected to the mainland as recently as 13,000 years ago (Des Lauriers, 2006). Despite this, the pollinator communities of Cedros Island are suggested to be depauperate when compared to the mainland (Brown & Faulkner, 1988). While the petals of 13770 are uniformly yellow with small central petal lobe red spots, those of CI have orange to red stripes on a yellow background that vary considerably among flowers, even within the same individual (Figure 1).

The ecological, geographic and abiotic contexts of floral evolution in section *Erythranthe* have received much less attention, despite ongoing work into the evolutionary ecology of *M. cardinalis* in response to recent drought and climate change (Angert et al., 2018; Sheth & Angert, 2014, 2016; Wooliver et al., 2020, Vtipil & Sheth, 2020; Coughlin et al., 2022). We are still untangling the effects of changing climate on floral traits and the pollinators that interact with and influence their evolution. When compared to leaf morphology and other physiological traits, floral traits remain understudied in the context of climate (Miladin et al., 2022; Caruso et al., 2003). Here, we employ a comprehensive integrative investigation of a suite of floral traits relevant to pollination from accessions of *Mimulus cardinalis* collected from seed across a latitudinal gradient of the species wide distribution in western North America. We relate observed patterns of floral trait variation to patterns of genetic and climate variation across the distribution of *M. cardinalis*, to better understand floral trait divergence in a geographic and environmental context. We highlight two distinct floral morphs from the northern and southern range extents which display a shift from red to yellow floral colour, to explore geographic and climatic correlates of floral transitions and potential pollinator shifts on a genetic level.

In doing so, we address the following research questions: How is variation in climate variables associated with the latitudinal range of *M. cardinalis*? What is the quantifiable degree of variation in pollinator-relevant floral traits across range-wide accessions of *M. cardinalis*, when grown in common greenhouse conditions? Is there more variation in some types of pollinator-relevant traits than others? How is observed floral trait variation potentially perceived by pollinators? Are there geographic gradients in these variable traits, or is geographic variation seemingly random? What are the possible genetic components associated with floral trait divergence at the range edges?

## MATERIALS AND METHODS

### Plant Material

This study focused on the typically hummingbird-pollinated species *Mimulus cardinalis* Douglas ex Benth. (syn. *Erythranthe cardinalis* (Douglas ex Benth.) Spach, Phrymaceae), a member of the developing model genus *Mimulus* section *Erythranthe* (Yuan 2019). Inbred lines CE10 and 13770 were obtained as described by (Wenzell & Neequaye *et al*., 2025). Red-flowered NFS084 was collected in southwest Oregon, USA (42.53311°N, - 123.77258°W). Red-flowered HOL012 was collected at Hollenbeck Canyon, California, USA, (32.67553°N, -116.826°W). These lines were provided by the Sheth Lab, North Carolina State University, USA. CI was collected from Cedros Island, Mexico, Baja California, and provided by Yaowu Yuan, University of Connecticut, USA. The reference lines 13770 and CE10 were inbred for at least 10 generations, whilst CI was inbred for an unknown number of generations and NFS084 and HOL012 were wild-collected seeds. Plants were grown from seed in common garden conditions in glasshouses at the John Innes Centre, Norwich, UK. Seeds were sterilised and grown as described by (Wenzell & Neequaye *et al*., 2025).

### Climate Data

Climatic conditions in the region of each *Mimulus cardinalis* line were extracted from the WorldClim 2.1 current datasets of environmental variables. WorldClim 2.1 current climate data is a monthly average for the years 1970-2000 (Fick & Hijmans, 2017).

Datasets for minimum temperature (°C), maximum temperature (°C), average temperature (°C), precipitation (mm), solar radiation kJ m-2 day-1), wind speed (m s-1), and water vapour pressure (kPa) at 30 second resolution (∼1 km^2^) were downloaded from WorldClim. These datasets were imported into R (version 4.3.2) using the R package ‘raster’ v3.6 for working with GeoTiff files (Hijmans, 2025).

A data frame containing the name of each *Mimulus cardinalis* line and coordinates for the population of each line was made in R. Whist accurate sampling origins were available for most of the lines, collection locality information for lines 13770 and CI were imprecise (based on locality descriptions only) and therefore don’t have reliably recorded exact coordinates. A closest best estimate was used for these data based on locality descriptions. Climatic conditions for each of the lines were extracted from the environmental variables datasets using the coordinates (known or inferred) for each line.

Boxplots were created using the R package ‘ggplot2’ v3.5.1 (Wickham, 2016). An ANOVA was used to compare the climatic conditions of each line and a Tukey’s HSD post-hoc test was used to identify which *Mimulus cardinalis* lines experience significantly different climatic conditions.

Principal Component Analysis (PCA) was performed in R to reduce WorldClim climate data to three principal components which were then plotted against each other.

### Existing Data

Data from lines CE10 and 13770 were originally published in (Wenzell & Neequaye et al., 2025) and not re-collected. Data presented here from CI, NFS084 and HOL012 are all original to this work.

### Morphology

Floral morphology was measured as described by (Wenzell & Neequaye *et al*., 2025). In brief, three flowers each from 5-15 individuals were measured with 150mm digital callipers (Linear Tools, RS components). Corolla length (base of corolla to mouth opening), display height (vertical spread of petal lobes, viewed from the front) display width (horizontal spread of petal lobes, viewed from the front), opening height (vertical space within corolla tube mouth), opening width (horizontal space in mouth) and herkogamy (difference between length of pistil from receptacle to stigma and length of stamens from receptacle to anthers) were measured to the nearest 0.1mm.

### Nectar Properties

Nectar properties were determined as described by (Wenzell & Neequaye *et al*., 2025). In brief, at least three flowers each of 5-15 individuals were sampled on the first day of opening; as the glasshouses are pollinator-free, this should represent the total nectar standing crop on the first day. Nectar for lines CE10, 13770, and NFS084 was collected into microcapillary tubes (Cat. No. 2930210, 1.55mm external diameter, Paul Marienfield GlmbH & Co. KG, Germany) from the corolla base. Height of nectar in tubes was measured in mm using digital callipers and converted into microlitres using capillary tube dimensions and the geometric formula for cylinder volume [height of nectar in mm/100*103.87=h/100*(π*0.5752)]. For lines CI and HOL012, a micropipette was instead used to remove nectar, which was removed in increments of 2μl until the corolla base was empty. Total nectar sugar concentration was measured to the nearest 1% using a nectar refractometer (0 to 50% (BRIX) sugar, Bellingham and Stanley UK Ltd).

### Floral Pigments

Pigments were extracted from whole corollas (1 flower per plant), including anthers but excluding stigmas and calyces. Fresh corollas had been flash-frozen in liquid nitrogen and stored at -80 degrees Celsius until use. Anthocyanins were extracted using methods modified from (Butelli et al., 2008). In brief, individual frozen corollas were ground in a pestle and mortar cooled with liquid nitrogen. 5mL of acidified methanol (0.3% HCl, vol/vol) was added to each sample. Samples were left to shake for 24 hours at 4 degrees Celsius. Samples were centrifuged at 10,000 rpm for 5 minutes.

Carotenoids were extracted using methods modified from (Sérino et al., 2009). In brief, corollas were ground in a pestle and mortar cooled with liquid nitrogen, 500 microlitres saturated NaCl solution and 250 microlitres of hexane were added, and the samples were vortexed at maximum speed for 30 seconds. 5mL ethyl acetate was added and samples were again vortexed at maximum speed for 30 seconds. The yellow upper organic phase was transferred to an Eppendorf tube and centrifuged at 10,000rpm for 5 minutes.

To determine total anthocyanin or carotenoid content, 500 microlitres of sample was transferred to a UV-transparent cuvette. To quantify total pigment, absorbance was measured at 525nm and 450nm for anthocyanins and carotenoids respectively, using a spectrophotometer (DS-11 FX UV-Vis-Spectrophotometer, Cambridge Bioscience, UK). For identification of individual anthocyanin and carotenoid compounds, 500 microlitres of sample were transferred to amber High Performance Liquid Chromatography (HPLC) vials and liquid chromatography mass spectrometry was performed (Agilent G6546A LC/Q-ToF, Santa Clara, CA, United States) as described in (Wenzell, Neequaye et al 2025). Compounds were identified using their retention times and mass spectra, and quantified using the equation [area under curve x 1000 / (mg tissue x molecular weight)] to give mass of compound in micrograms/milligram of corolla tissue (as in Kellenberger et al., 2019).

### Reflectance Spectrophotometry and Perception Modelling

Reflectance spectrophotometry and perception modelling as described in (Wenzell & Neequaye et al., 2025). Floral color was quantified by measuring reflectance of fresh corolla tissues (collected immediately prior from plants growing in the glasshouse) using a reflectance spectrometer (FLAME-S-UV-VIS-ES Assembly, 200–850 nm, Ocean Insight), with light source from a pulsed xenon lamp (220 Hz, 220–750 nm, Ocean Insight, v. 2.0.8) on four areas of the corolla: the upper petal lobe, the central (bottom) petal lobe, the lower side petal lobe, and the throat of the corolla tube, including any nectar guide. The reflectance spectrometer collected readings every 5 ms and averaged 25 scans per reading (Ocean View spectroscopy software, Ocean Insight). Readings were standardized with an absolute white color reference (Certified Reflectance Standard, Labsphere) and electric dark was used as the black standard. Three readings (technical replicates) were taken from each tissue of each flower measured, and three flowers (biological replicates) were measured per individual plant, with 5-16 plants characterized per line. Reflectance curves at wavelengths from 300 to 700 nm were visualized for all four tissues. All three petal lobe tissues showed similar reflectance curves, so we present and analyze only the central petal lobe tissue hereafter, as this petal lobe is forward facing (not strongly reflexed) and thus visible to pollinators approaching the front of the flowers.

Floral color (reflectance) was compared among lines using a principal component analysis (PCA) using R v.4.2.287, package psych version - 2.2.592 to allow for the varimax rotation, which prioritizes loading each trait (i.e., wavelength) onto only one PC axis, based on the intensity of reflectance at a given wavelength (based on intervals of every ten nanometers from 300 to 700 nm, which captures wavelengths relevant to the vision of most pollinators). First, a PCA was run based on intensity values at all 41 wavelengths; then, only factors with an eigenvalue>1 were used in the PCA, which resulted in a final PCA of 3 factors. To visualize whether these flowers reflect in the ultraviolet (UV) range (including potential patterning), which is important for bee vision, we photographed flowers of 1–3 individuals of all four color morphs under UV light using a UV-sensitive camera (Nikon D610, converted to a Full Spectrum camera by removing the Kolari Vision UV bandpass filter, with a Micro-Nikkor 105 mm lens) in a dark room illuminated by a UV black light.

To characterize how floral colors may be perceived by trichromatic bee pollinators, we plotted the reflectance data of lower central petal lobes on trichromatic models of bee (*Apis mellifera*) visual systems using R v.4.2.287 with R package pavo version 2.7.1109, which estimates level of contrast of a color signal against a vegetative (green) background and displays the results on the bee colour perception hexagon. In addition, we used the pavo package to characterize how floral colours may be perceived by tetrachromat pollinators (including birds and day-flying butterflies) and visualized these in a three-dimensional tetrahedral space with the vertices representing the four cone receptors.

### Floral Scent Collection

Floral scent collection was performed as described in (Wenzell & Neequaye et al., 2025). Floral Volatile Organic Compounds (VOCs) were collected for 3 technical replicates of at least 5-10 plants per line (grown in the John Innes Centre glasshouse), largely from the same individuals that were phenotyped for morphology, nectar, and pigment. Floral VOCs were collected through dynamic headspace collection, as described by (Wenzell & Neequaye *et al*., 2025). Pairs of freshly cut fully open flowers (or no flowers, but otherwise identical, for a blank/empty control) were placed in 50 mL glass beakers containing approximately 40 mL of sterile ddH2O, which were placed in PTFE-lined oven roasting bags (Sainsbury’s, UK), sealed and connected to scent traps with aluminium twist ties. Scent traps were composed of modified glass Pasteur pipettes filled with 100mg Porapak Q (Merck/Sigma), contained within pea-sized amounts of silanized glass wool on either side, and were connected via silicone tubing to Spectrex PAS-500 volatile pumps (Merck/Sigma), which were run at a flow rate of 100 mL/min for 24 hours to capture any circadian variation in emissions. Floral VOCs were then eluted from scent traps using 600 uL of extraction solvent (10% HPLC-grade acetone in ≥ 95% HPLC-grade hexane) into 2mL screw-top amber glass vials, which were stored at -20°C in a dedicated scent freezer before concentration and GCMS analysis.

Compound GC-MS analysis— Sample aliquots of 150ul were first concentrated under ambient room air to 50ul prior to injection of 3ul into an Agilent GC-MS system (7890B GC with 5977A/B MS) using a Gerstel MPS autosampler system with a splitless inlet held at 250C. The oven temperature was held at 50C for 4 minutes, then raised at 5C/minute to 230C, where it was held for 4 minutes. The column was a Phenomenex 7HG-6015-02-GGA (35m x 250um x 0.1um) and the carrier gas was helium at a flow rate of 1.0603 ml/minute. The MS was run in scan mode for ion masses between 50 and 600, with the MS source held at 230C.

### Floral Scent Analysis

Results were analysed using Agilent Unknowns Software (v10.1), generating peaks and offering the top three potential NIST library identifications. Peaks with areas less than 1 x 10^5^ were excluded due to the sensitivity of the Unknowns integrator software. Peaks eluting after 30 minutes are unlikely to be VOCs and were excluded. Sample peaks with the same library hit and retention time of a compound in the blank, and with area less than 5-fold higher than the blank peak were excluded. Peaks where all three hits contained silica or phthalate contaminants were also excluded. Mass spectra of the remaining uncertain peaks were visually compared to the blank sample’s mass spectra at similar retention times and retained or excluded based on the criteria above. After combining results from all samples of a given line, peaks were retained if they were present in at least ⅔ of the samples from that line, and discarded otherwise.

For further verification of peaks, an alkane ladder (Merck, 49452-U) was run at the same time as samples and the Kovats Retention Index (KRI) was calculated and compared with published NIST values.

Compounds with an available standard were verified through GCMS and quantified by converting peak areas to ng/flower/hour. Compounds without an available standard but with matching spectra and KRI to published NIST data were quantified using a standard with a similar structure. Compounds without matching spectra and KRI could not be verified and therefore were left as ‘Unknowns’ and quantified using standards with a similar structure to the top hit from the NIST library. Pie charts were plotted using the R(Team, 2021) ‘pie’ function. Radius was scaled in accordance with the total emission per line. Non-metric Multidimensional Scaling (NMDS) was carried out with the R (R Core Team, 2021) package ‘vegan’ (Oksanen, 2010) using Bray distances.

### Whole Genome Analysis

Whole genome analysis was as performed in (Wenzell & Neequaye et al., 2025). DNA was extracted from leaf tissue of CI using the NUCLEON PHYTOPURE 50 PREPX0.1G Kit (Cytiva Catalogue ID: RPN8510) with the addition of an RNAse treatment using ThermoScientific RNAase T1 Catalogue #EN0541. Library preparation and sequencing was performed by Novogene using short-read Illumina sequencing technologies. All genome and transcriptome data are available on ENA, with accession number PRJEB98459. Reads were aligned to the publicly available *Mimulus cardinalis* CE10 reference genomes (CE10g_v2.0) (http://mimubase.org/) using bwa version 0.7.17 (Li et al., 2009a) with subsequent processing using samtools version 1.7 (Li et al., 2009b) and bcftools version 1.10.2 (Li, 2011).

### Gene Expression Analysis

RNA from inbred lines CI, 13770, and CE10 (excluding wild-collected NFS084 and HOL012) was extracted from whole corollas, including anthers but removing calyx, ovary, and stigma tissue, such that it was comparable to pigment analyses. This was performed using the Spectrum^TM^ Plant Total RNA Kit (Merck Catalogue #STRN50-1KT) and samples were eluted in water. Three replicates per genotype were sent for sequencing by Novogene. Messenger RNA was purified from total RNA using poly-T oligo-attached magnetic beads. Following this, it was fragmented, and first strand cDNA synthesis was performed using random hexamer primers. This was followed by second strand cDNA synthesis. End repair, A-tailing, adapter ligation, size selection and amplification and purification were then performed to complete the RNA library. Kallisto (https://pachterlab.github.io/kallisto/)(Bray et al., 2016) was used to align reads from *M.cardinalis* to the *M.verbenaceus* reference genome (MvBLg_v2.0, Mimubase) (Wegryzn and Yuan, 2023) to produce gene read counts, as this genome had the highest gene-level assembly quality of the *Mimulus* genomes available.

Transcriptome data from 13770 and CE10 was published in (Wenzell & Neequaye et al., 2025).

### Statistical Analyses

Statistical significance of differences in morphology, nectar properties, pigment quantities, total scent emission and gene expression were performed using one-way ANOVA. Where there were significant differences between lines, Tukey’s Honestly Significant Difference (HSD) with a 95% confidence interval was used to determine which pairwise comparisons were significantly different. These analyses were performed using the R(Team, 2021) version 4.3.2 functions ‘aov’ and ‘Tukey HSD’. The NMDS plot and ANOSIM analyses were performed using vegan v2.6-4. Bar plots were plotted using ggplot2 v3.4.4 and geom_beeswarm v0.7.2.

## RESULTS

### Mimulus cardinalis populations experience different climatic conditions across a latitudinal range

The species *M. cardinalis* occurs in a latitudinal gradient across western North America, spanning from central Oregon, USA to northern Baja California, Mexico (Angert et al., 2018) (Figure 1B). To gain insight into the ecological conditions that may drive floral trait diversification in *M. cardinalis*, we analysed local climate data from a 30-year period (1970-2000) from the sampling locations of each of the following accessions occurring across the species range; NFS084, 13770, CE10, HOL012 and CI (pictured in Figure 1A).

Climate data shows the variation in climatic conditions experienced by these accessions across their range, with significant variation found in all climate variables tested, except for solar radiation (Figure 1, Supplementary Figure S1). Maximum and minimum temperatures were found to be significantly different at the five accession sampling locations (F=3.545, p=0.0121, F=19.84, p<0.0001 for maximum and minimum respectively) (Figure 1C, Figure S1A). Minimum temperature was not significantly different between the northern and central lines NFS084, 13770 and CE10, but was significantly different when the northern and central lines were compared to the more southern HOL012 and CI (p < 0.01 for all 3 comparisons to HOL012, p < 0.001 for all 3 comparisons to CI, respectively) (Figure 1C).

This trend is consistent with average temperature (F=7.265, p=0.0000902), which is not significantly different between NFS084, 13770 and CE10 but was significantly higher for HOL012 (p < 0.05 for all 3 comparisons) and higher still for CI (p < 0.05 for all 3 comparisons) (Figure 1D). As with temperature, precipitation was also significantly different (F=9.296, p<0.0001) where the northernmost line, NFS084 experiences the most precipitation (p<0.05 when compared to 13770, HOL012 and CI), whilst CI experiences the least (p<0.0001 when compared to NFS084) (Figure 1E). Consistent with increased temperature and reduced precipitation at the southern range edges, water vapour pressure was also significantly different between locations (F=25.35, p<0.0001) where the same trend as that of temperature is observed (Figure 1F). There is no significant difference between the northern and central locations but significantly higher water vapour pressure in the case of the southern accessions HOL012 and CI (p<0.05, p<0.001 for HOL012 and CI respectively). We also found a significant difference in wind speed (F=13.5, p<0.0001) but not in solar radiation (F=1.237, p=0.306) (Supplementary Figure 1B, 1C, respectively). It is important to note that these populations occur at variable elevations, which are not accounted for separately here. Wind speed and solar radiation are important factors when considering variable elevations, which is known to influence local climate and ecological factors (Oldfather et al., 2020).

### Floral morphology of M. cardinalis varies across the species range

Variable ecological conditions, including climate, are known to interact with floral traits in *M. cardinalis* (Sheth & Angert, 2014, 2016; Vtipil &Sheth, 2020). Using climate as a proxy marker for abiotic variation, we wanted to quantify the degree of floral trait variation in *M. cardinalis* accessions from across the geographic range. Floral morphology has a direct impact on pollinator attraction and effective pollination. Morphological changes in floral structures are frequently associated with transitions in pollination strategy (Whittall & Hodges 2007). To understand the intraspecific variation in floral morphology of *M. cardinalis* across its species range, corolla length (CL), display height (DH), display width (DW), opening height (OH) and opening width (OW) were measured for the accessions NFS, 13770, CE10, HOL012 and CI (Figure 2A).

**Figure 2.**
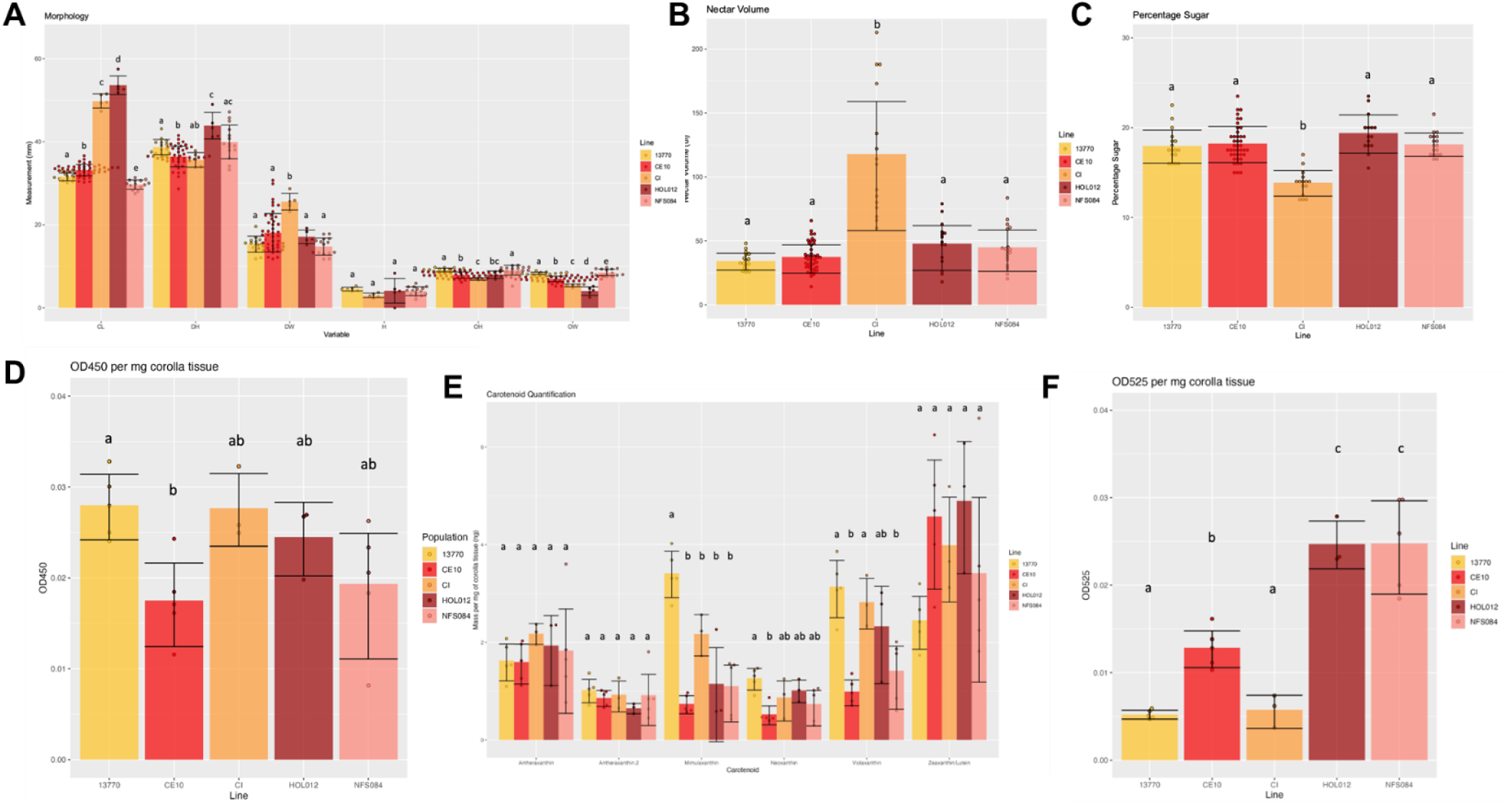
Floral Trait Variation. (A) Morphological measurements of the *Mimulus cardinalis* accessions CE10, HOL012, NFS, CI and 13770. CL = corolla length, DH = display height, DW = display width, H = herkogamy, OH = opening height, OW = opening width. Bars show the mean measurement, and error bars indicate mean ± standard deviation. (B) the nectar volume in microlitres per flower and (C) percentage sugar in nectar. (D) Absorbance at 450nm, indicating total quantity of carotenoids, and (E) quantities of individual carotenoids. Numbered compounds indicate a different isomer. (F) Absorption at 525nm, indicating total quantity of anthocyanins. Significance values were obtained using one-way ANOVA, followed by Tukey’s HSD for pairwise comparison. Bars sharing a letter are not significantly different, and lettering is independent for each variable. Bars show mean expression, while error bars represent mean ± standard deviation.

Corolla length was significantly different between all five accessions, with the southern line HOL012 having the longest average corolla length, followed by the southern CI (F=498.9, p<2e-16). Similarly, HOL012 has the greatest display height (F=12.49, p=2.11E-08), whilst CI has the greatest display width (F=9.735, p=1.76E-06). In contrast, CI has the smallest opening height (F=14.06, p=9.35E-09) whilst all accessions were variable in opening width (F=68.33, p<2e-16).

The northernmost accessions, 13770 and NFS084, had significantly shorter corolla lengths and significantly greater opening heights and widths than any of the other accessions (F=498.9, p<2e-16), supporting previous findings where 13770 was compared only with CE10 (Wenzell & Neequaye et al., 2025). Herkogamy was also measured but no significant differences were found between accessions.

### Cedros Island M. cardinalis produces large quantities of nectar

Alongside morphological transitions, nectar production also plays a key role in pollinator choice in *Mimulus* section *Erythranthe* (Schemske & Bradshaw 1999). Nectar volume (Fig. 2B) was significantly greater in CI than in all other accessions (F=49.3, p=<2e-16). In contrast, percentage total sugar (Fig. 2C) was significantly reduced in CI relative to all other accessions (F=25.16, p=9.59E-12). CI therefore produces a higher volume of more dilute nectar than the other four studied *M. cardinalis* accessions.

### Carotenoid profiles of M. cardinalis accessions show limited variability

Carotenoids are responsible for yellow floral pigmentation in *M. cardinalis* corollas (LaFountain et al., 2015). Carotenoid pigments were extracted from whole corollas and absorbance at 450nm was measured (Fig. 2D) to quantify total carotenoids. The only significant difference in total carotenoids of the givr accessions was found between the yellow accession 13770 and red-coloured accession CE10 (p = 0.03), as was reported in (Wenzell & Neequaye et al., 2025). Individual carotenoid compounds were identified by UHPLC-MS and their quantities followed the same trend. Individual carotenoids were not significantly different across accessions, except for violaxanthin (p=0.00033), neoxanthin (p=0.007) and mimulaxanthin (p<0.0001), which were significantly higher in 13770 than CE10, as reported in (Wenzell & Neequaye et al., 2025) (Fig. 2E).

### M. cardinalis accessions each produce a unique profile of anthocyanins

Anthocyanins are responsible for red pigmentation in *Mimulus* (Pollock et al., 1967), including the variable red corolla stripes and spots characteristic to CI and 13770 respectively. In 13770, they were shown to be significantly reduced relative to the red line CE10 (Wenzell & Neequaye et al., 2025). Anthocyanins were extracted from flowers of CI and the previously uncharacterised red-flowered HOL012 and NFS084, and total anthocyanins were quantified by measuring absorbance at 525nm (Fig. 2F). Total anthocyanins were significantly lower in the yellow-flowered accessions 13770 and CI when compared to the three red accessions (F= 38.3, p<0.00001). The previously uncharacterised accessions HOL012 and NFS084 show significantly higher total anthocyanins than range-centre accession CE10 (p = 0.0008, p=0.0002 for HOL012 and NFS084 respectively).

To get a better understanding of the diversification of anthocyanin biosynthesis *M. cardinalis*, individual anthocyanins were characterised and quantified using UHPLC-MS. Presence or absence of different compounds is shown in Table 1. Anthocyanin biosynthesis was found to be highly variable, with no two lines sharing the same anthocyanin profile in terms of individual anthocyanin presence or absence. Moreover, of those anthocyanins that are shared between two accessions, all were found to be significantly different in their individual quantitative amounts, except for an isomer of pelargonidin-glucoside-rhamnose (Figure S2).

**Table 1.** Presence (green) and absence of individual anthocyanin isomers in 5 accessions of *M. cardinalis*.

|  | 13770 | CE10 | CI | HOL012 | NFS084 |
| --- | --- | --- | --- | --- | --- |
| Pel-Glc 1 | N | N | N | Y | Y |
| Pel-Glc 2 | N | N | Y | Y | N |
| Cyan-Glc | Y | Y | N | N | N |
| Pel-GlcRha 1 | Y | Y | N | N | Y |
| Pel-GlcRha 2 | N | Y | Y | Y | Y |
| Cyan-GlcRha | Y | Y | N | N | Y |
| Pel-Glc-Ac 1 | N | N | N | Y | N |
| Pel-Glc-Ac 2 | N | N | Y | Y | N |
| Cyan-Glc-Ac 1 | N | N | Y | N | N |

### The yellow morphs at range-edges are genetically distinct from the range-centre red morph

In this study, we focused genetic analyses on the two yellow-coloured populations at the furthest range edges, the northern range-edge 13770 and southern range-edge CI. When comparing whole genomes of CI and 13770 to the range-centre accession CE10, CI was found to have greater genetic variation compared to CE10 than the same comparison between CE10 and 13770. This is according to the scoring of whole genome alignments, in which CI is found to have greater singletons and lower percentage mapping and pairing to the CE10 genome, when compared to 13770 (Table S1).

To further investigate how genetic variation may relate to the floral traits we quantified, we looked at specific examples of genes potentially associated with pollinator-relevant floral traits. Genes potentially involved in nectar production were identified based on their functional annotation in the *Mimulus verbenaceus* genome (MvBLg_v2.0, Mimubase) (Wegryzn and Yuan, 2023). Their expression in CI, 13770 and CE10 was analysed using transcriptomic data from these lines (Fig. 3A). One of these genes encodes a putative bifunctional monodehydroascorbate reductase and carbonic anhydrase, corresponding to *Nectarin III* (*NEC3*) in *Nicotiana benthamiana* which is thought to buffer nectar pH and support nectar oxidative balance (Carter and Thornburg, 2004). This gene showed significantly higher expression in CI than in either of the other lines (F=18.25, p=0.00281).

**Figure 3.**
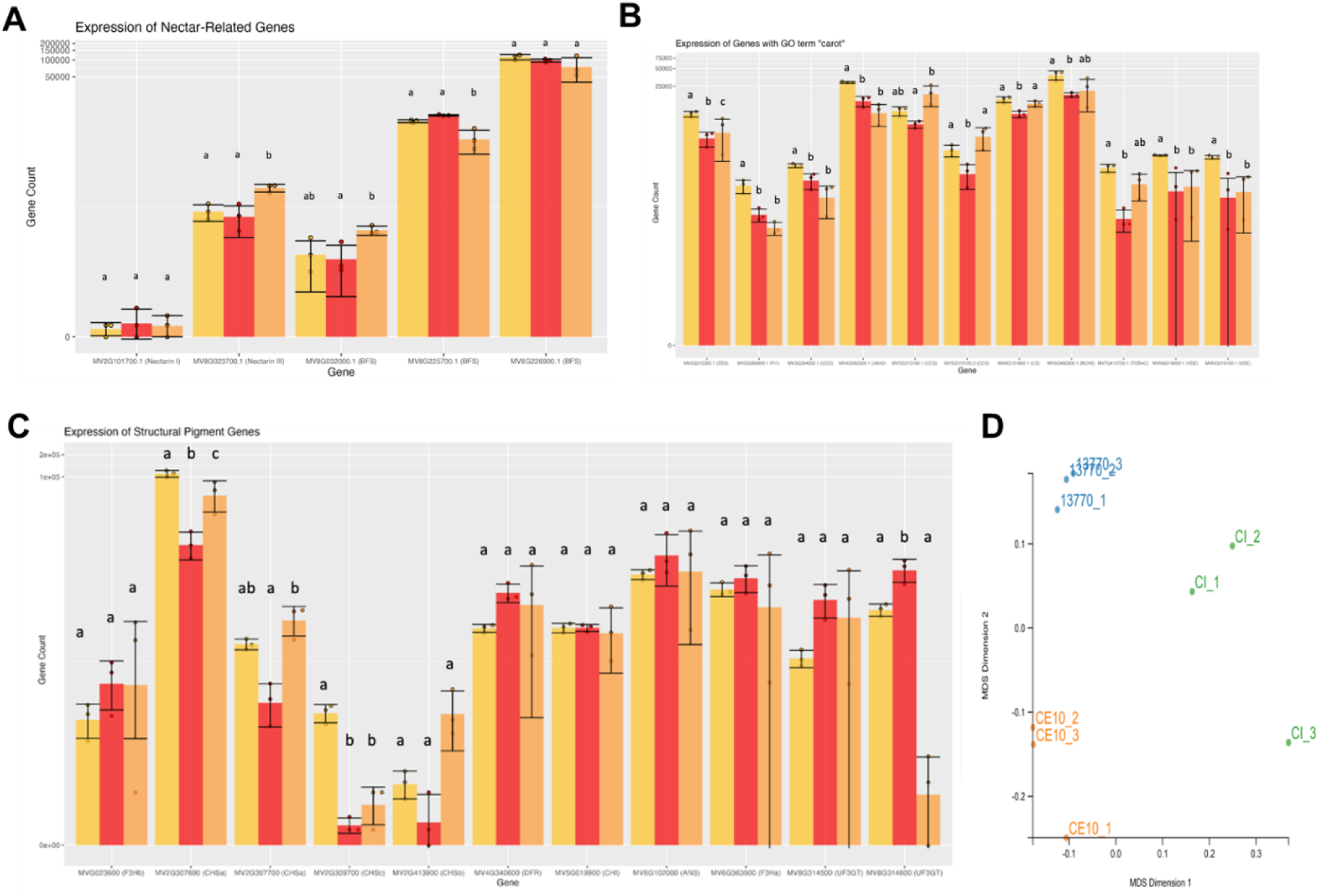
Gene Expression. (A) Expression of genes involved in nectar production in the *M. cardinalis* accessions CE10, CI and 13770. (B) Expression of significantly differentially expressed genes annotated with the GO term ‘carot’. (C) Expression of genes encoding structural enzymes in the anthocyanin biosynthesis pathway. The y-axis is displayed as a logarithmic scale to account for large differences in expression. Bars show the mean measurement, and error bars indicate mean ± standard deviation. Statistical significance was determined using one-way ANOVA followed by Tukey’s HSD. Bars that share letters are not significantly different from one another. (D) MDS of whole transcriptome data generated in Degust using edgeR.

Additionally, differentially expressed genes found using the GO terms ‘nectar’ and ‘nectar secretion’, included three putative beta-fructofuranosidases (*BFS*es) which hydrolyse sucrose to produce D-Fructose and D-Glucose (Chen et al., 2021). One of these copies had significantly higher expression in CI compared to CE10 (F=6.89, p=0.0279), and another had significantly lower expression in CI than in either of the other lines (F=26.41, p=0.00106) (Figure 3A). There was no significant difference between lines in the third (F=1.842, p=0.238). The increased expression of *NEC3* and a homolog of *BFS*, and the downregulation of another homolog of *BFS*, correlate with the increased nectar production found in CI, which is higher than any other *M. cardinalis* accession characterised to date. Meanwhile, beta-fructofuranosidases hydrolyse sucrose (Heil et al., 2005) and are implicated in maintaining sugar balance in extrafloral nectar in multiple species (Nogueira et al., 2018). These genes are not necessarily the cause of nectar differences in CI but may play a role in maintaining important nectar parameters across the larger nectar volume found in this line.

Genes involved in carotenoid biosynthesis were found to have varying differences in expression in CI relative to 13770 and CE10 (Fig. 3B). Notably, the genes with statistically significant differences in expression all encode enzymes directly involved in carotenoid biosynthesis. In CI, only two genes were significantly upregulated relative to CE10, and none were significantly more greatly expressed in CI than 13770. Both up-regulated genes compared to CE10 were putative copies of *CCS*, which catalyses the conversion of antheraxanthin and violaxanthin into capsanthin and capsorubin, respectively (Camara & Moneger 1981). In contrast, seven genes were upregulated in 13770 relative to CE10, three of which were also significantly elevated relative to CI, which are involved in various steps of carotenoid biosynthesis and catabolism. 13770 therefore seems to have more widespread upregulation of carotenoid biosynthesis than CI, relative to CE10. This is consistent with its increased carotenoid content as published in (Wenzell & Neequaye et al., 2025).

Genes encoding structural enzymes in the anthocyanin biosynthesis pathway have previously been identified in *Mimulus lewisii*, the sister species of *M. cardinalis* (Yuan et al., 2014). Expression of putative homologs of *Chalcone Synthase a (CHSa)*, *CHSb, CHSc, Chalcone Isomerase (CHI), Flavonoid 3-Hydroxylase a (F3Ha), F3Hb, Dihydroflavonol 4-reductase (DFR), Anthocyanidin synthase (ANS)* and *UDP-3-O-glucosyltranserase (UF3GT)* (Yuan et al., 2014) were compared between the three sequenced lines (Fig. 3C). Expression of two *CHSa* homologs was significantly greater in CI than in CE10, and one of these was also significantly greater in CI than 13770 (F=23.39, p=0.0122). Relative to CE10, both CI and 13770 lines show overall upregulation of *CHS* expression. Expression of one *MlUF3GT* homolog was significantly reduced in both CI and 13770, relative to CE10 (F=19.43, p=0.00239). In all other cases, structural anthocyanin biosynthesis genes were not significantly greater in red CE10 compared with the two yellow lines.

We also looked at potential anthocyanin regulators (Figure S3). *ANbHLH1* expression was significantly higher than CE10 in both yellow lines, and significantly higher in CI than 13770 (F=28.32, p=0.000878) (Supplementary Figure S3A). Homologs of *WD40a, b* and *c* genes were also identified and their expression quantified (Supplementary Figure S3B).

Expression of a *WD40b* homolog was significantly greater in both yellow lines than in CE10 (F=10.4, p=0.0112). Expression of a *WD40c* homolog was significantly greater in CI than in either 13770 or CE10 (F=47.52, p=0.000209). These encode proteins that form part of a MYB-bHLH-WD40 (MBW) complex that co-ordinately activates at least four (F3H, DFR, ANS, UF3GT) enzymes of the anthocyanin biosynthetic pathway (Yuan et al., 2014).

Some spatially restricted R2R3-MYBs controlling anthocyanin production, *Petal Lobe Anthocyanin (PELAN)* and *Nectar Guide Anthocyanin (NEGAN)* (Yuan et al., 2014, LaFountain et al., 2023), positively regulate anthocyanin biosynthesis in *Mimulus* section *Erythranthe* petal lobes and nectar guides respectively. The expression of putative *PELAN* homologs was compared (Supplementary Figure S3C). Two of the genes were not expressed at detectable levels in CI and 13770, while they were expressed in CE10, consistent with suggestions that *PELAN* expression is absent in yellow *M. cardinalis* (Yuan et al., 2014, Wenzell & Neequaye et al., 2025). In all cases where expression of putative *PELAN* homologs differed between yellow lines, it was greater in CI than in 13770. This is in agreement with the intermediate anthocyanin profile of CI when compared to red-flowered CE10 and yellow-flowered 13770. A single homolog of *MlNEGAN*(Yuan et al., 2014) was identified and its expression quantified (Supplementary Figure S3D). This gene was expressed significantly more in 13770 than in CI or CE10, consistent with the red throat spotting found in 13770 only (F=62.29, p=0.000097).

The overall pattern of variation in transcriptomes reflects that of the whole genomes, in which the two yellow-flowered range-edge accessions are genetically distinct from the range-centre CE10 accession as well as from each other (Figure 3D). We hypothesize that genetic variation at the genomic and transcriptomic level in *M. cardinalis* is related to the variation in floral traits observed and will be influenced both by abiotic factors such as climate and biotic factors such as pollinator interactions.

### Floral Pigment Perception by Pollinators Shows Variability in M. cardinalis

Following the characterisation of the genetics and chemistry of floral pigment variation in accessions of *M. cardinalis*, we sought to identify how pigments are perceived. Floral pigments play a pivotal role in pollinator attraction and are often associated with transitions in pollination syndromes (Bradshaw & Schemske 2003). In order to characterise the in-situ tissue pigmentation patterns of *M. cardinalis*, reflectance of a series of four areas on the inflorescence were measured (Figure 4). In contrast with all other known *Mimulus cardinalis* lines, CI produces striped flowers, providing an almost intermediary pigment pattern when compared to its red (CE10, HOL012, NFS084) or yellow (13770) counterparts. The upper side petal lobe (Fig. 4A,E) and lower side petal lobe (Fig. 4B,F) of CI was found to overlap entirely with its yellow counterpart 13770, showing a distinct contrast with all other red-flowered lines. The lower central petal lobe (Fig. 4.C,G) has a more distinctive reflectance pattern, differentiating across accessions. The corolla throat shows the greatest overlap of all *Mimulus cardinalis* accessions as all accessions share their characteristic red throat-spotting (Figure 1A).

**Figure 4.**
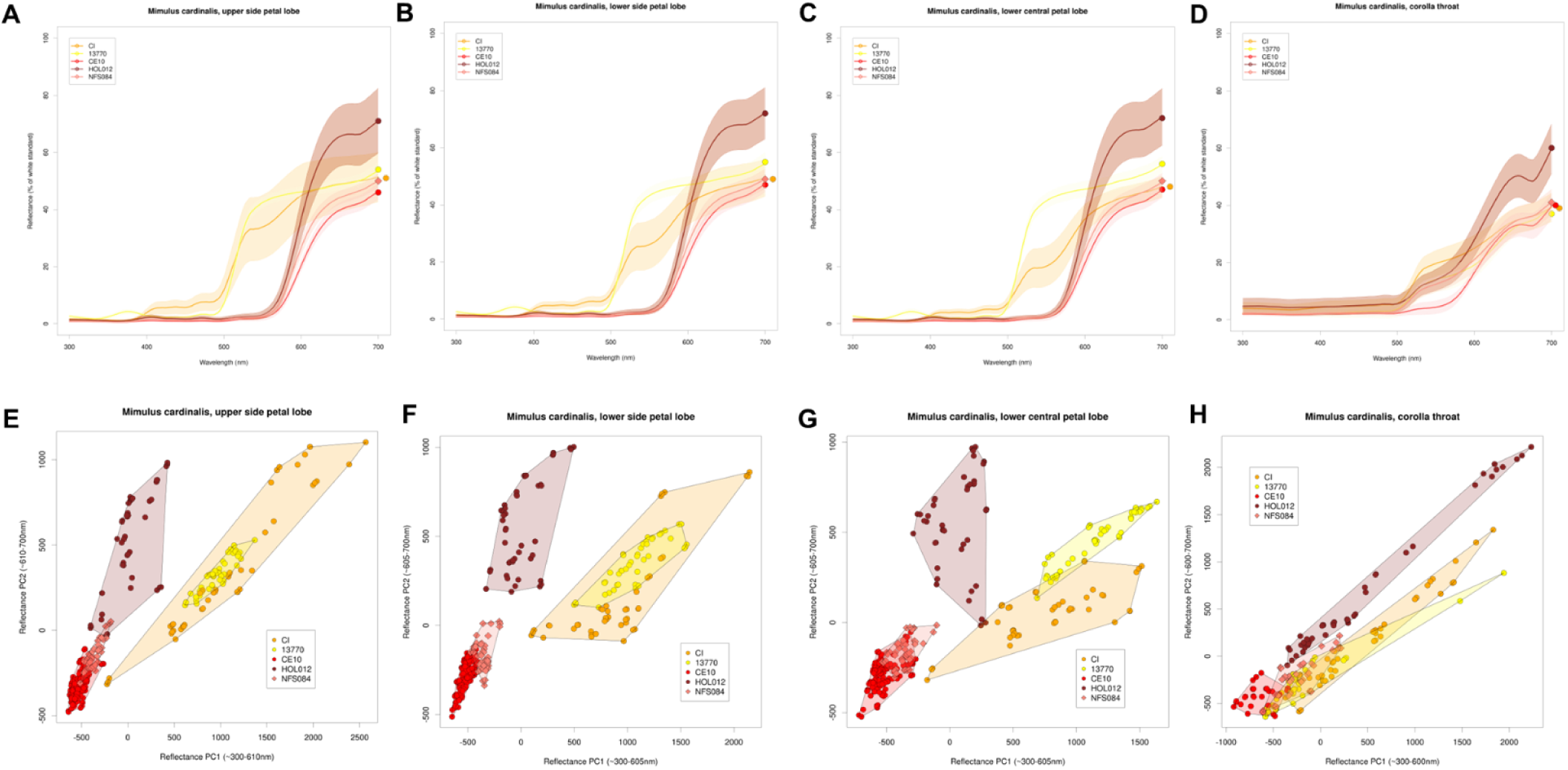
Reflectance. Spectral reflectance of (A) upper side petal lobes (B) lower side petal lobes (C) lower central petal lobes (D) corolla throat spots and of M. cardinalis lines. Principal component analysis of spectral reflectance values of of (E) upper side petal lobes (F) lower side petal lobes (G) lower central petal lobes (G) corolla throat spots and of M. cardinalis lines red lines CE10, HOL012 and NFS and yellow lines 13770 and NFS.

We then sought to model reflectance to known perception models of two distinct insect pollinator guilds. Models of bee perception are based on the trichromat honeybee, *Apis mellifera*, and rely on contrast from a “less perceived” achromatic centre, where distance from the centre suggests higher perceptibility. The yellow-flowered accessions CI and 13770 more are more perceivable by bees than their red counterparts, with distinct perceptibility separating the two yellow accessions. The yellow accessions do not overlap in tissues studied, except for the characteristic, red-spotted corolla throat where all accessions are equally less perceptible than the central petal lobe of the yellow lines (Fig.5A-D).

**Figure 5.**
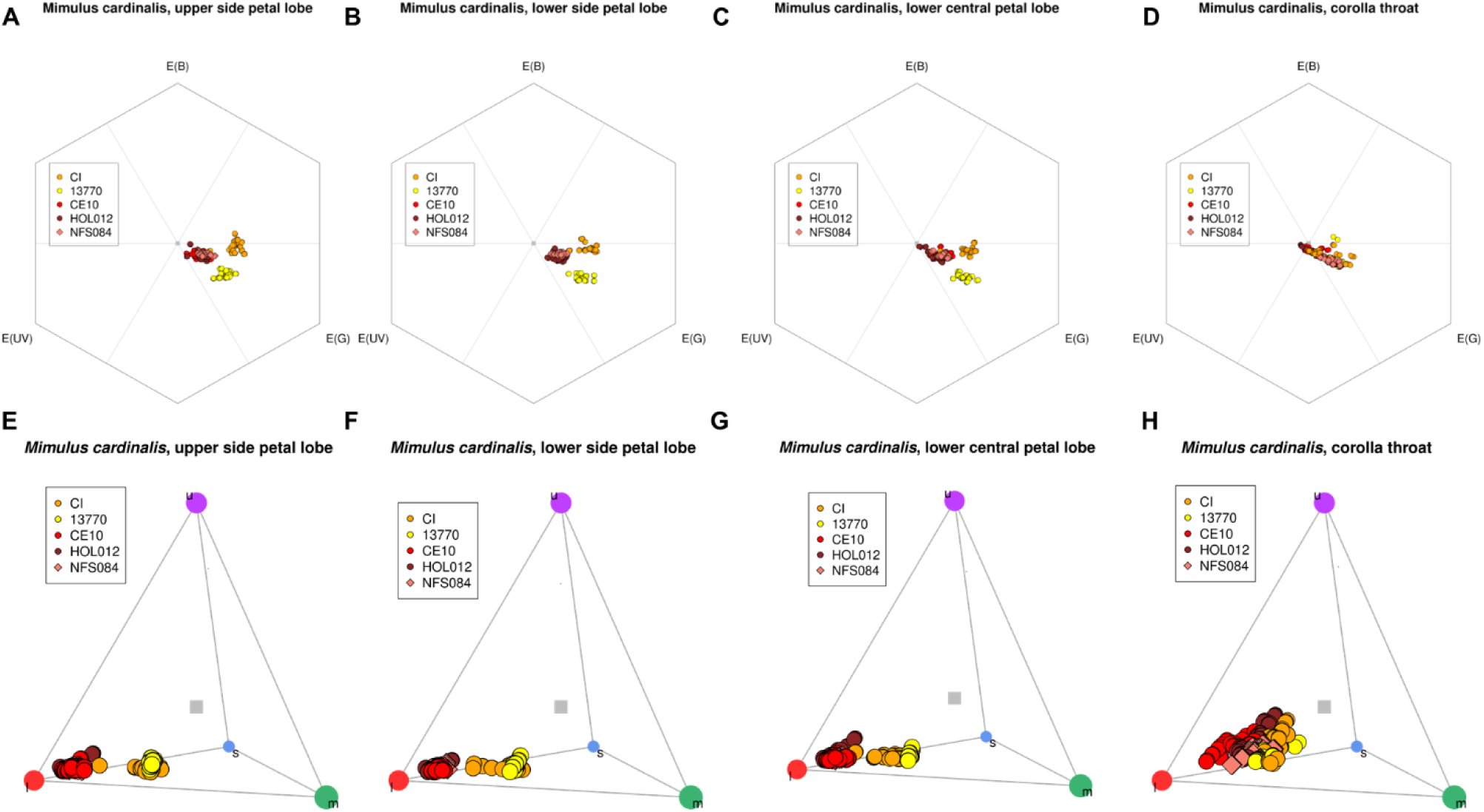
Perception. Bee perception model based on spectral reflectance of (A) upper side petal lobes (B) lower side petal lobes (C) lower central petal lobes (D) corolla throat spots and of *M. cardinalis* lines. Diurnal lepidopteran and hummingbird perception tetrachromat model based on spectral reflectance of (E) upper side petal lobes (F) lower side petal lobes (G) lower central petal lobes (H) corolla throat spots and of *M. cardinalis* accessions.

Models of perception by day-flying avian and lepidopteran pollinators, such as hummingbirds and butterflies, show a similar pattern of perception, in which CI and 13770 show separation from the red-flowered accessions, which cluster together (Figure 5E-F). It is worth noting that red-striped CI accessions were used in the generation of the reflectance data perception models.

### Cedros Island M. cardinalis produces high volumes of floral scent

Floral scent plays an important role in pollinator perception and attraction (Raguso, 2008). The floral volatile organic compounds (VOCs) produced by CI, HOL012, and NFS084 were captured using dynamic headspace collection and compared to previously published data from 13770 and CE10 (Wenzell & Neequaye et al., 2025). All accessions were found to have a unique floral scent profile (Figure 6, Table S2).

A total of 57 VOCs were detected across all *M. cardinalis* accessions, with no volatile found in more than 3 accessions. A total of 7 compounds were found in 3 lines each (Unknown KRI=949, Lavender lactone, (*Z*)-Linalool oxide, Methyl salicylate, Hexyl-2-methylbutyrate, Phenethyl-2-methylbutyrate, and Unknown KRI=1685). A total of 9 compounds were found in 2 lines each ((α)-Pinene, Sabinene, (β)-Pinene, 1-Octen-3-ol, (*Z*)-Arbusculone, (*E*)-Linalool oxide, Monoterpenoid KRI=1096, Unknown KRI=1272, and Valencene). The remaining 41 compounds were found to be unique to one line or another. Volatile profiles are significantly different between lines overall (ANOSIM: R= 0.956, p=0.001), and even amongst the 16 shared compounds, no compounds share the same levels of production between lines.

Accessions show variation in the relative proportions of VOCs, at the individual VOC and class of VOC level (Fig. 6A). The similarity of scent profiles between lines was analysed using non-metric multidimensional scaling (NMDS; Fig. 6B) where all accessions clustered distinctly, particularly CI. HOL012 was found to have the highest total volatile emissions (Figure 6C), particularly in aromatic and terpenoid volatiles (Figure 6D and E, respectively) (overall ANOVA: F_4_ = 92.3, p < 2e-16; Tukey HSD for HOL012 *versus* other lines: p < 0.0001 for all comparisons). CE10 produces lower volatile emissions of fatty-acid derivatives (overall ANOVA: F_4_ = 17.52, p = 2.5e-10; Tukey HSD for CE10 *versus* other lines: p < 0.001 for all comparisons). CI scent contains the largest proportion of non-aromatic, non-FAD, non-terpenoid, non-unknown volatiles (Figure 6G). CI also produces a lower proportion of methyl salicylate than other lines and a similar proportion of lavender lactone when compared to the range-centre accession CE10. Meanwhile, 13770 has a higher proportion of lavender lactone compared with other lines, but lacks methyl salicylate (Wenzell & Neequaye et al., 2025).

**Figure 6.**
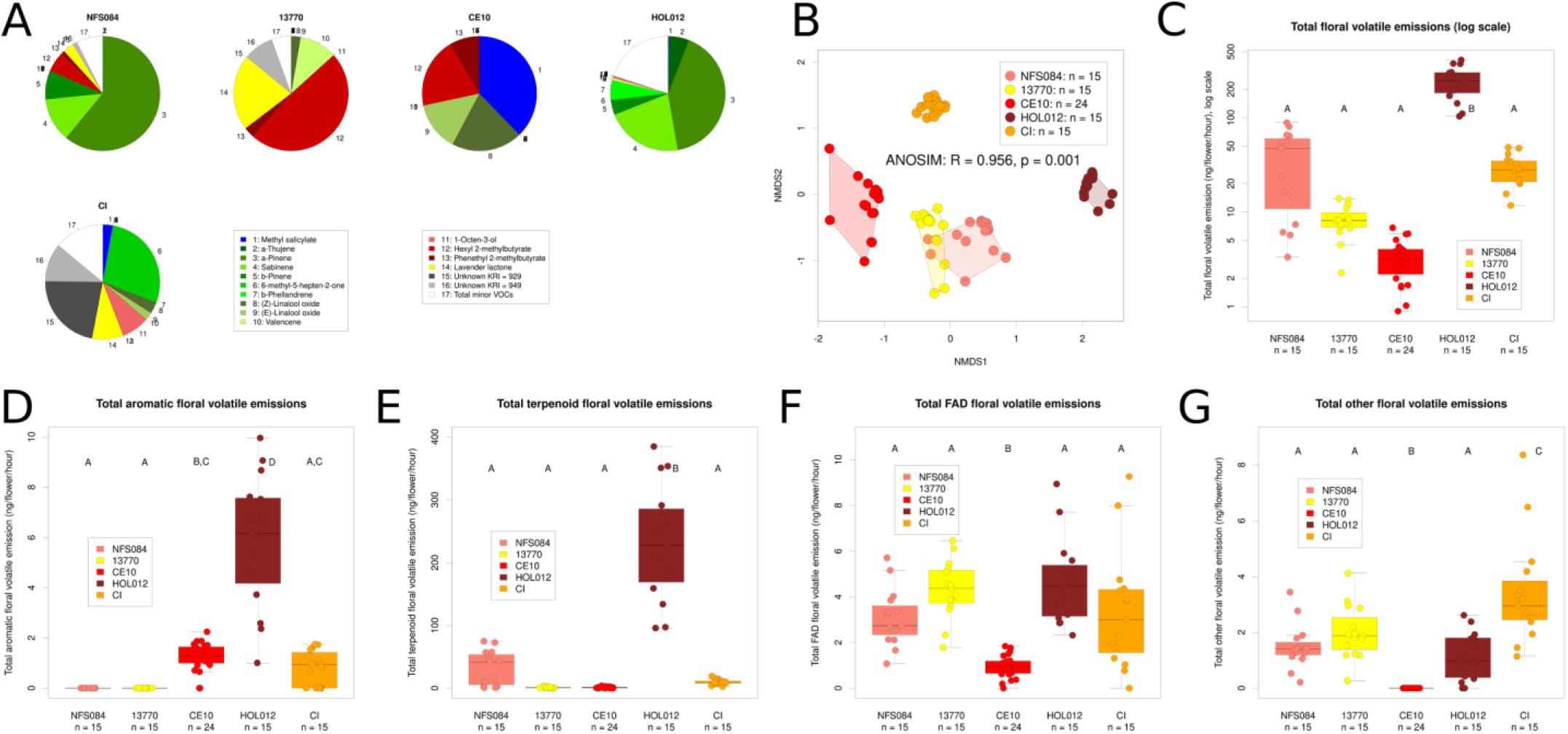
Scent. (A) Proportions of total scent of individual VOCs as pie charts. Colour indicates chemical class; green = terpenoid, red = lactone, yellow = fatty acid derivative, blue = aromatic, grey = unknown. (B) NMDS of all volatiles across accessions (C) Total volatile emissions (log scale) per flower per hour (D) Total aromatic VOCs per flower per hour (E) Total terpenoid VOCs per flower per hour (F) Total FAD VOCs per flower per hour (G) Total VOCs not belonging to the 3 main classes (D-F) (excluding unknowns). Significance values were obtained using one-way ANOVA, followed by Tukey’s HSD for pairwise comparisons. Bars sharing a letter are not significantly different, and lettering is independent for each variable. Bars show mean expression, while error bars represent mean ± standard deviation.

## DISCUSSION

We characterized a suite of floral traits in accessions from across the *Mimulus cardinalis* species range, finding a high level of intraspecific variation in multiple traits in this theoretically phenotypically coherent species. We found the highest variability in scent and anthocyanin production (including extreme disparity in individual anthocyanin and volatile profiles), with carotenoids and morphology seemingly more stable between accessions. The variation appears to be inconsistently distributed across the geographic range and therefore range of climatic factors, which are known to directly and indirectly influence pollinator-relevant floral traits but may play a lesser role in this system given the lack of geographic trends in most floral traits (with the exception of some morphological traits).

### Range-wide floral trait variation and pollinator perception in M. cardinalis

We found high variability in floral traits of geographically widespread accessions of *M. cardinalis*. The two accessions found towards the southern range-edge, CI, and to a lesser extent HOL012, were both found to have significantly greater corolla lengths than the other, more northern accessions. For HOL012 this morphological variation also included greater display height, whilst CI had great display width. Floral morphological changes in the southern range edge line CI did not show the same pattern of differences from the red-flowered line CE10 as the independently derived yellow-flowered 13770 line from the northern range edge (Wenzell & Neequaye et al., 2025). This lack of convergence is consistent with comparisons to another red to yellow shift in the closely related species *M. verbenaceus,* also in section *Erythranthe*, with the yellow line MvYL having a longer corolla, a larger opening and no significant difference in display width relative to the conspecific red line MvBL (Wenzell & Neequaye et al., 2025). In other words, the three derived yellow lines do not display consistent morphological shifts in comparison to their conspecific red lines.

The shorter corolla and wider opening of 13770, in comparison to CI and other red-flowered *M. cardinalis* lines could favour pollinators such as bees, which struggle to access nectar rewards in flowers with long, narrow corollas, which they have even been observed to tear open (Giovanetti et al., 2023; Wenzell & Neequaye et al., 2025). In contrast, narrow-billed hummingbirds or pollinators like hawkmoths - which use a long proboscis to access nectar and do not typically land - may not be hindered by these parameters and would therefore be effective pollinators for CI (Grant & Temeles, 1992; Aigner & Scott, 2002). The significant increase in display width and height in CI and HOL012, respectively, may also influence pollinator attraction. For example, increased floral size has been shown to contribute to hawkmoth preference in *Jasminum fruticans* (Thompson, 2001) and *Nicotiana* species (Kaczorowski et al., 2012). Meanwhile, hummingbird pollinated flowers are often smaller (Kaczorowski et al., 2012).

The range-edge accession, CI, was also found to produce significantly higher volumes of more dilute nectar than any other *M. cardinalis* line studied here, with more dilute nectar, perhaps as a trade-off to maintain an equal production of costly sugars. Changes in nectar properties may influence pollinator behaviour, as large volumes of dilute nectar are associated with hummingbird pollination (Lanza et al., 1995, Dalsgaard et al., 2009, Kay and Grossenbacher, 2022) and have been suggested to be adapted to hawkmoth pollination as well (Wang et al., 2023). Conversely, many bee-pollinated flowers have reduced nectar volume and higher sugar concentration (Martínez-Díaz et al., 2023). Nectar composition may also influence these interactions, as observed in *Nicotiana* species, where specific nectar sugars and amino acids are strongly correlated with pollinator preference (Tiedge and Lohaus, 2017). Studies have found a discordance between where hummingbird-pollinated plants occur and where hummingbirds themselves occur, instead finding that hummingbirds will visit flowers when conditions are adequate to make hummingbird attraction and reward trait values, for example through nectar rewards (Coffey & Simons, 2025; Ramírez-Burbano et al., 2022).

In addition to nectar rewards, floral pigments play a key role in mediating plant-pollinator interactions (Weiss, 1991). Individual patterns of carotenoid accumulation across accessions reflects patterns of accumulation between red and yellow morphs to that observed in (Wenzell & Neequaye et al., 2025). In contrast, anthocyanin pigment biosynthesis shows high levels of variation across *M. cardinalis* accessions in this study. Of the five accessions studied, no two accessions produced the same anthocyanin profile, even within the red-flowered CE10, NFS084 and HOL012. Interestingly, HOL012 is the only accession with only pelargonidin (no cyanidin) as its core anthocyanidin, which may contribute to its distinction in the reflectance in the PCA space, despite appearing as the same red as NFS084 and CE10 to the human eye. This reflects the highly branched pathway of anthocyanin biosynthesis, in which numerous anthocyanin decorations can be added to the same anthocyanidin core or different anthocyanidin cores can be used (Saigo et al., 2020). Whilst carotenoids have a linear pathway of production, reducing the opportunities for significant deviations, anthocyanin profiles can change in species through the up or down-regulation of single enzymes (Rausher et al., 1999). This is reflected in the expression of genes associated with anthocyanin production, which show greater variation/deviation between accessions in this study than those involved in carotenoid biosynthesis.

In addition to scent perception, pigments pay a key role in pollinator perception. Different pollinators have different visual systems and hence perceive colour differently. Hummingbirds can perceive a wide colour spectrum and typically prefer red or white flowers which have high contrast against a green vegetative background using their tetrachromatic visual system (de Camargo et al., 2019). The red pigments of HOL012, NFS084 and CE10 therefore lend themselves well to hummingbird perception (and the use of red can act to exclude bees, Castellanos et al., 2004) yet still use different suite of anthocyanins amongst themselves to produce the similarly perceived red corollas. In contrast to hummingbirds, models of bee colour perception showed that yellow *Mimulus* flowers had greater contrast against the vegetative background than their red conspecifics (Wenzell & Neequaye et al., 2025), and many bee species lack a red receptor (Peitsch et al., 1992). This was seen in CI, where floral reflectance showed a greater similarity to yellow-flowered 13770 than to its red-flowered counterparts in terms of its ability to be perceived by bees, but was still distinct from 13770 with no overlap of perception. Similarly, hawkmoths (as represented by *Manduca sexta*) have been shown to prefer *M. lewisii x M. cardinalis crosses* with mutations producing paler yellow, white or pink flowers, as opposed to red (Byers and Bradshaw, 2021) and they show a strong preference for yellow *M. aurantiacus* over red in the wild (Streisfeld & Kohn, 2007). It is worth noting that the work by Streisfeld & Kohn (2007) in *M. aurantiacus* takes place in and around southern California, overlapping with the southern range extent of *M cardinalis*. We therefore suggest that this could be consistent with hawkmoths as important potential pollinators in this area, especially as hawkmoths are known to be involved in shifts from hummingbird pollination in *Mimulus* (Byers & Bradshaw, 2021; Streisfeld & Kohn, 2007). It is also notable that both southern accessions CI and HOL012 had increased corolla length relative to all other *M. cardinalis* accessions in this study, which is consistent with expectations for hawkmoth pollination (Johnson et al., 2017; Alexandersson & Johnson, 2002).

Patterning is also an important component in pollinator attraction. Bee-pollinated flowers typically have a nectar guide (de Camargo et al., 2019), and bees had more trouble handling yellow *M. verbenaceus,* which lacks a nectar guide, than they did *Mimulus cardinalis* line 13770, which has red spots in the nectar guide area (Wenzell & Neequaye et al., 2025). Petal lobe patterning can also be important in addition to nectar guides. For example, hawkmoths have been suggested to have an innate preference for radially patterned flowers (Kelber, 2002), and moth pollinated *Gladiolus* species often have pale, tubular flowers, with darker, mottled patterns (Goldblatt and Manning, 2002). As hummingbirds are still known to visit the yellow-flowered morph of *M. cardinalis* 13770 (Vickery & Vickery 1992), this pigment transition in CI to yellow flowers, which may be more attractive to insect pollinators, may represent a reproductive assurance mechanism to cope with declining hummingbird populations while attracting a novel and potentially more reliable pollinator in addition. Rather than representing an “anti-bird” adaptive walk, CI may instead be transitioning to a more generalist “pro-insect” strategy (Castellanos et al., 2004) in an environment that may be depauperate in potential pollinators (Brown & Faulkner, 1988).

As with anthocyanins, scent production is highly biochemically variable across accessions of *M. cardinalis*. Most noteworthy is an increased total VOC production, more specifically higher aromatic and terpenoid volatiles, in HOL012, an accession found towards the southern range-edge. Not only are total levels of VOC production highly variable between accessions, but the specific changes in scent composition are also highly variable, with each accession producing a unique bouquet of floral VOCs. Most hummingbird-pollinated flowers (including *M. cardinalis*, Byers et al. 2014a) are weakly scented (Knudsen et al., 2004) owing to hummingbirds having poor scent perception and memory (Goldsmith & Goldsmith 1982, Núñez et al., 2021). In contrast, floral scent is crucial in determining the behaviour of insect pollinators (Byers et al., 2014a,b; Byers, 2021). This existing high level of volatile emissions in accessions such as HOL012 may act as a “bridge” to enable a switch to other pollinators or a more generalised pollination syndrome.

Different pollinators also respond to specific chemicals, influencing floral scent profile. For example, *Clarkia breweri* is the only member of its genus that is moth pollinated, and it produces high quantities of linalool oxide (Raguso and Pichersky, 1995) and methyl salicylate, the latter inducing a strong electroantennographic response in the hawkmoths *Sphinx perelegans* (Raguso and Light, 1998) and *Manduca sexta* (Hoballah et al., 2005). Linalool oxide is found in the scent composition of range edge accessions CI and 13770. Different pollinators can also have different responses to the same chemical, for example, methyl salicylate. Methyl salicylate is found in southern edge accessions CI and HOL012 and is the most predominant compound in the VOC composition of range centred accession CE10. This compound is suggested to be attractive to hawkmoths (Hoballah et al., 2005, Raguso and Light, 1998) as well as a bee repellent (Henning et al., 1992, Mayer, 1997).

### Genetic variation at range edges in M. cardinalis

Genetically, populations at range edges are expected to have variable (either higher or lower) genetic variation than populations found in more central areas of the geographic species range (Ehrlén & Morris, 2015; Pulliam, 2000; Sagarin et al., 2006). We found the northern range-edge accession 13770 and southern range-edge accession CI to be genetically distinct at the whole genome and transcriptome level, in comparison to the range-centre accession CE10. Southern range-edge accession CI was found to share less whole genome sequence homology and patterns of floral-trait gene expression compared with CE10 than 13770 in whole genome and transcriptome analyses. Coughlin et al., 2022 propose that this standing genotypic variation in *M. cardinalis* has allowed these populations to occupy a range of ecological niches and that this variation may serve to “buffer” *M. cardinalis* from environmental change. This genetic variation in relation to niche occupation is also explored by (Preston et al., 2022) who use transcriptome analyses to propose that rapid adaptation to climate variability in *M. cardinalis* is a result of high phenotypic plasticity. Meanwhile, (Vtipil & Sheth, 2020) suggest *M. cardinalis* persists across its wide geographic range due to plastic shifts in phenology, evolution of drought tolerance and movement to track climatically suitable habitats. As a result, *M. cardinalis* may be able to exploit both significant standing genetic variation as well as high levels of phenotypic plasticity when adapting to novel environments.

We found variation in expression of genes associated with floral traits, including pigments, which we hypothesize to confer the variation in floral traits observed in these accessions, as the three sequenced lines are inbred glasshouse-grown and not wild-collected lines. Because we used mainly inbred accessions in a common garden experiment, we propose that the variation observed in floral phenotypes is largely influenced by the genetic variation we found in the two yellow-flowered, range-edge populations. 13770 and CI represent not only novel floral phenotypes in their accumulation of pigments but in their sites of origin, in relation to the model, range-centre accession CE10.

The southernmost *M. cardinalis* populations have been found in previous studies to be the most responsive to short-term climate changes and to have the highest phenotypic plasticity in relation to genetic variation (Sheth & Angert, 2016; Wooliver et al., 2020). Our results are consistent with these findings as we find the highest genetic variation in CI when compared to a similarly pigmented northern 13770 yellow morph of *M. cardinalis*. Moreover, we find divergence in expression patterns of genes related to floral pigment phenotypes. All accessions studied, even those that don’t vary much in human-visible colour, such as the three red morphs (CE10, NF084 and HOL012), all produce distinct anthocyanin profiles. This is consistent with the high variability in expression of structural genes and regulators of this pathway found between the range-centre CE10, northern range edge 13770 and southern range edge CI. This high variability in expression of pathway genes suggests that anthocyanins are complicated in this group, even for lines we expected to be the same due to similar visible colour phenotypes. The variation in floral trait associated genes from across CI, alongside the climate data and historical studies, suggest a possible ecological adaptation to the pollinator-depauperate and climatically variable Cedros Island (Brown & Faulkner, 1988). “The Baldwin effect” describes the link between transcriptional plasticity and establishment of populations in new environments through novel adaptive trait emergence in addition to post-establishment range expansions (Baldwin, 1896; Levis & Pfennig, 2016; Santi et al., 2020). We hypothesise that standing genetic and phenotypic variation, as well as high levels of phenotypic plasticity, in *M. cardinalis* has allowed it to occupy this wide latitudinal range of niches, and that it has been able to differentiate novel floral traits to promote establishment in varied climatic niches.

### Floral trait variation in relation to climate

Floral trait macroevolution has a complex relationship with climate. Climate influences floral morphology directly and indirectly (Miladin et al., 2022). Climatic variables play a role in maintaining local pollinator communities, therefore influencing the pollinator-mediated selection of floral traits in plants within a geographic and therefore climatic range. For example, in South America, pollinator occurrence is known to be directly affected by numerous climate variables, including temperature (in the case of butterflies) though this is less consequential for hummingbirds, the predicted pollinator of *M. cardinalis* (Abrahamczyk et al., 2011). Further speciation following hummingbird specialisation is rare, likely due to the ability of the hummingbird to maintain gene flow, inhibiting population fragmentation as the hummingbird is a strong flyer and can cover an extensive range (Abrahamczyk & Renner, 2015). This is interesting in the context of *M. cardinalis* where local climate is highly distinct between sampling locations. In addition, as many hummingbird-related floral phenotypes such as increased carotenoid production and lack of scent seem due to loss-of-function mutations in *M. cardinalis* (*e.g.* Bradshaw & Schemske, 2003; Byers et al., 2014b), it is unclear how further novel phenotypes may emerge. More work is required to unpick the relationship between pollinator, climate and the traits that interact with both drivers of diversification/speciation.

Miladin et al., 2022 found vegetative traits, such as leaf size, to be directly influenced by climate, in addition to floral traits such as flower size. Other traits, including calyx shape, are directly influenced by pollinator identity and not climate, whereby the climatic influence on pollinators provides an indirect influence of climate on floral morphology (Miladin et al., 2022). Muir & Angert, 2017 show reduced plasticity in vegetative phenotypes of *M. cardinalis* in response to variable climatic conditions. Flowering time, a highly coordinated floral trait, shows plasticity in response to climate variables (Sheth & Angert, 2016; Vtipil & Sheth 2020). A recent study by (Albano et al., 2025) suggests that many *M. cardinalis* populations currently occupy climatic niches outside of their optimum conditions. In our study we also found high variability in pollinator-relevant floral traits in relation to the variable climates and ecological niches occupied by the five focal study accessions. It is likely that, despite existing across a latitudinal range, the various locations in which the geographically widespread accessions are established represent uniquely differentiated climatic and therefore ecological niches, as evidenced by the high variability found in all climate variables studied.

Many interacting and highly integrated physical, ecological, biotic and abiotic conditions and evolutionary processes determine the limits to where a species occurs, can occur or will occur in the future (Sexton et al., 2009). Local adaptation of populations across a range may drive individual populations to occupy different climate niche limits and therefore local ecologies. This is how climate change continues to shape range dynamics. It is important to consider the relationship between the climatic variables described here and the species range and how local topography, vegetation and elevation interact to drive variation in key ecological traits such as those required for effective pollination and outcrossing (Sexton et al., 2009, Oldfather et al., 2020). Studies such as those described here are important to consider in the context of changing global climate, particularly with a current megadrought occurring where many of the accessions are derived from, making the area the driest it has been for over a millennium (Williams et al., 2022).

## CONCLUSIONS

This study provides a comprehensive analysis of floral traits from five geographically widespread accessions of *Mimulus cardinalis*. We use genetic and climatic data to understand potential drivers of the diverse floral phenotypes observed across *M. cardinalis* accessions. Future work will directly measure biotic (pollinators) and abiotic (climate) factors *in situ*, providing in-depth population-level sampling, rather than use of the inbred lines as proxies used to facilitate this work. This work provides important context in the study of floral trait microevolution and the high levels of biochemical diversity that can be found within seemingly identical accessions of the same species. This work is especially important when considering conservation efforts and the managing of species in geographically diverse and rapidly changing climates.

## ACKNOWLEDGEMENTS

The authors thank Jay Goldberg and Pirita Paajanen for providing bioinformatic support during this project. The authors thank Seema Sheth and Sheth lab members for providing seeds for lines NFS084 and HOL012. Additionally, we thank Yaowu Yuan for seeds of CI and 13770 and H. D. Bradshaw, Jr. for seeds of CE10. Thanks also to Lionel Hill and Paul Brett for their aid with mass spectrometry.

