## Supplementary Materials for "In-depth phenotyping reveals unexpected floral trait variation in *Mimulus cardinalis* across a range-wide latitudinal gradient"

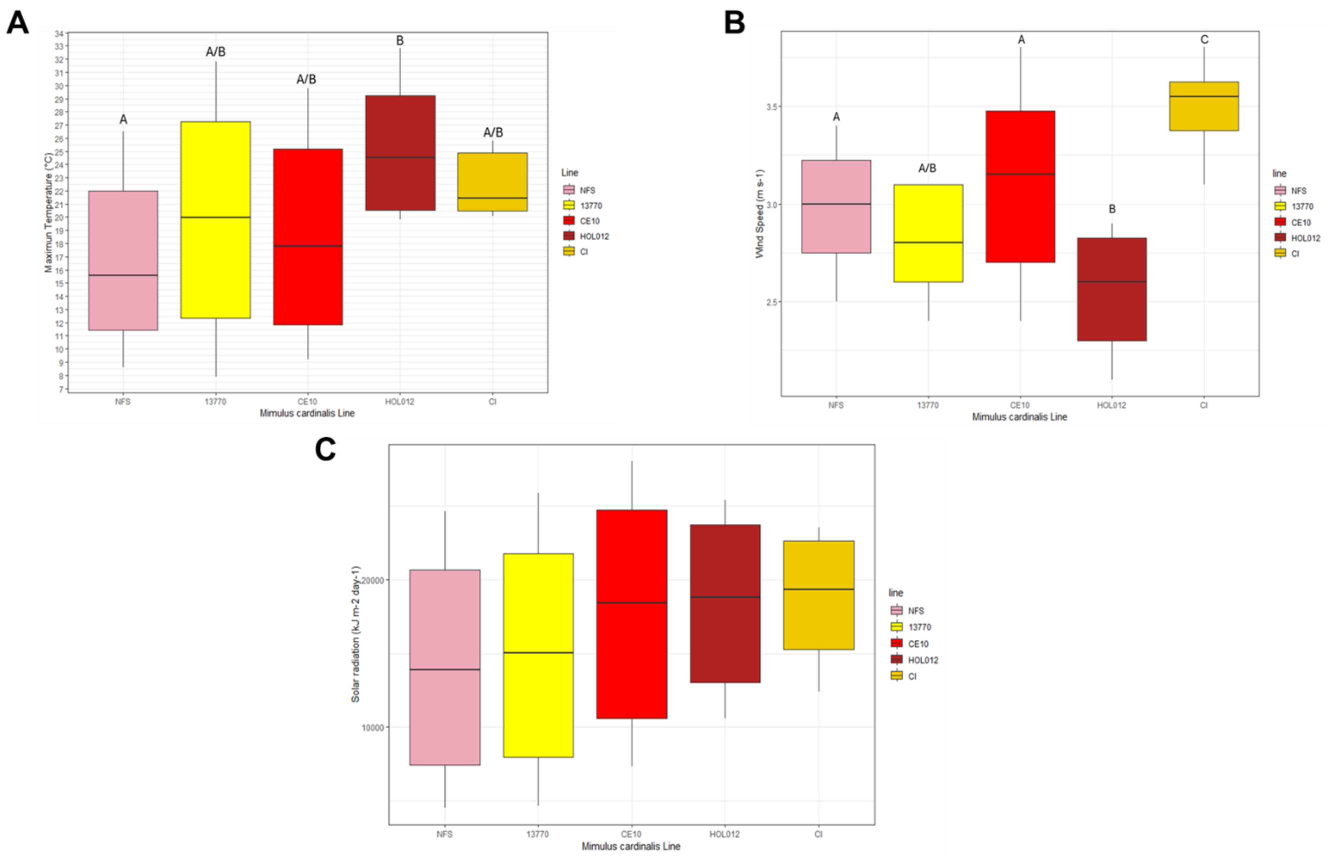


**Figure S1** - Climate variables (A) Maximum temperature (B) Wind speed (C) Solar radiation. For (A) and (B) bars sharing letters are not significantly different from one another, and lettering is independent for each variable. No bars for (C) solar radiation are significantly different from one another. Climate data is based on best estimates of seed sampling locations. Variance shows a 30-year average from 1970-2000. Significance values were obtained using one-way ANOVA, followed by Tukey’s HSD for pairwise comparison. Bars show median expression, while error bars represent mean ± standard deviation.

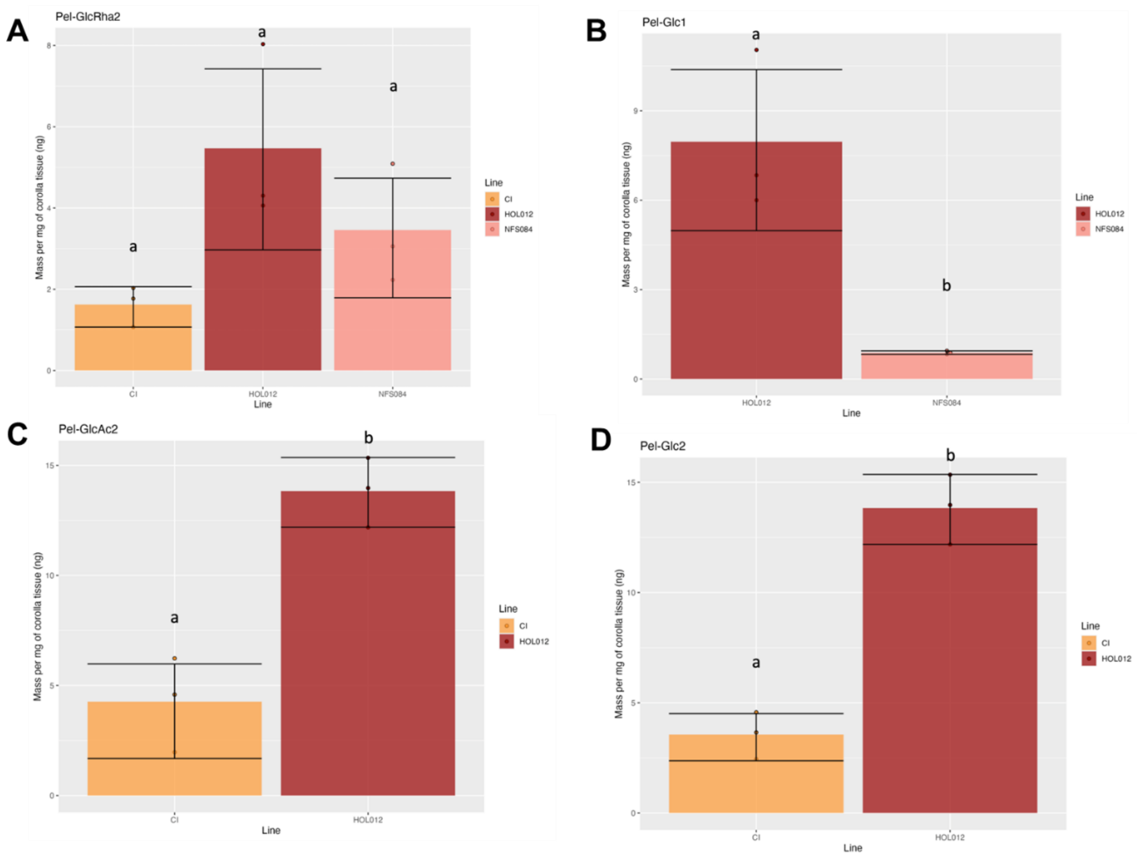


**Figure S2**- Quantification of Shared Individual Anthocyanins. Mass per mg of flower tissue (ng) of (A)Isomer 1 of Pelargonidin-glucoside-rhamnose (Pel-GlcRha2) in CI, NFS084 and HOL012. (B) Isomer 2 of Pelargonidin-glucoside (Pel-Glc2) (C) Isomer 2 of Pelargonidin-glucoside-acetate (Pel-Glc-Ac2) and (D) Isomer 2 of Pelargonidin-glucoside (Pel-Glc-2). Bars represent the mean value, and error bars represent mean ± standard deviation. Statistical significance was determined using one-way ANOVA, followed by Tukey's HSD.

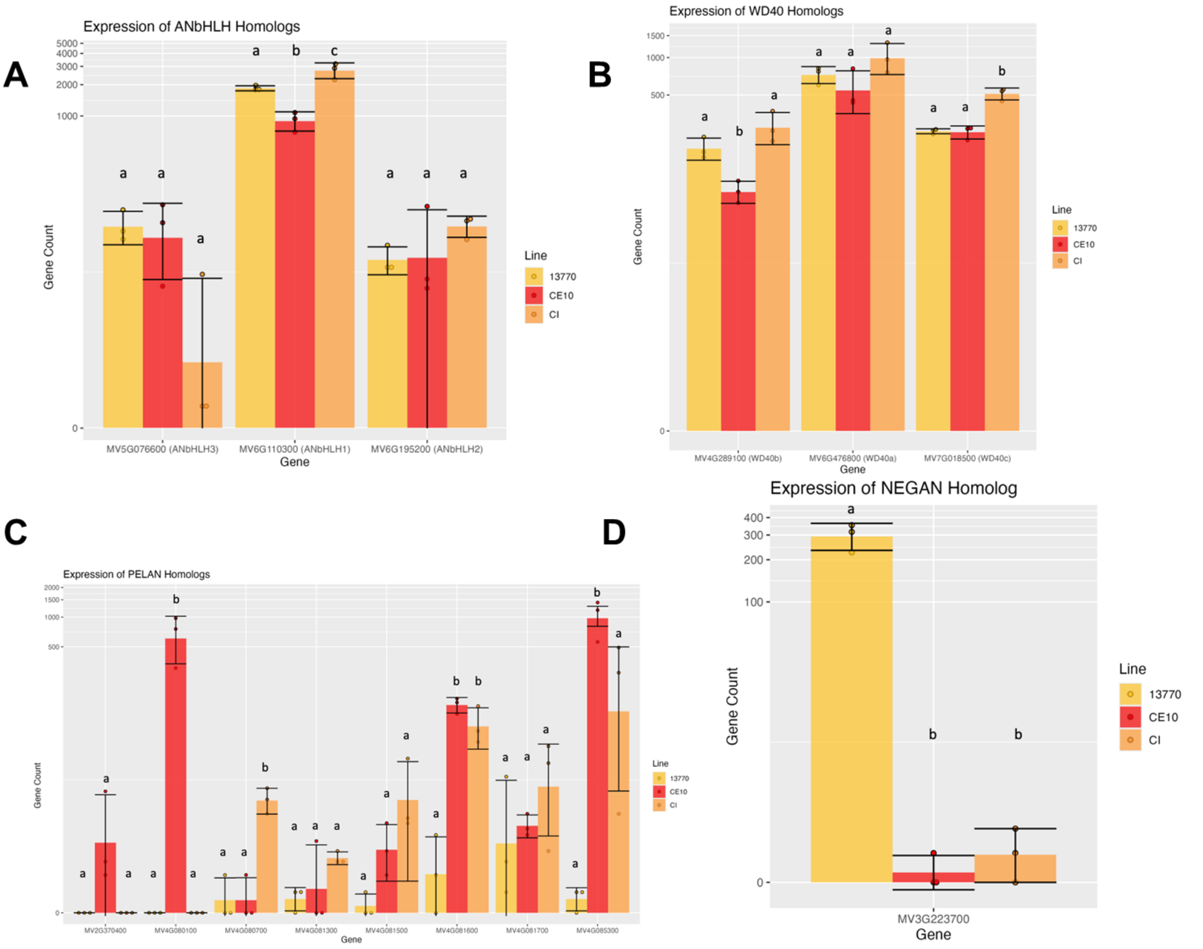


**Figure S3** – Expression of genes encoding MBW complex components: (A) ANbHLH homologs (B) WD40 homologs (C) PELAN homologs, (D) NEGAN homolog. Statistical significance was determined using one-way ANOVA, followed by Tukey's HSD. Bars sharing letters are not significantly different from one another and lettering is independent for each gene. The y-axes are displayed as a logarithmic scale to account for large expression differences.

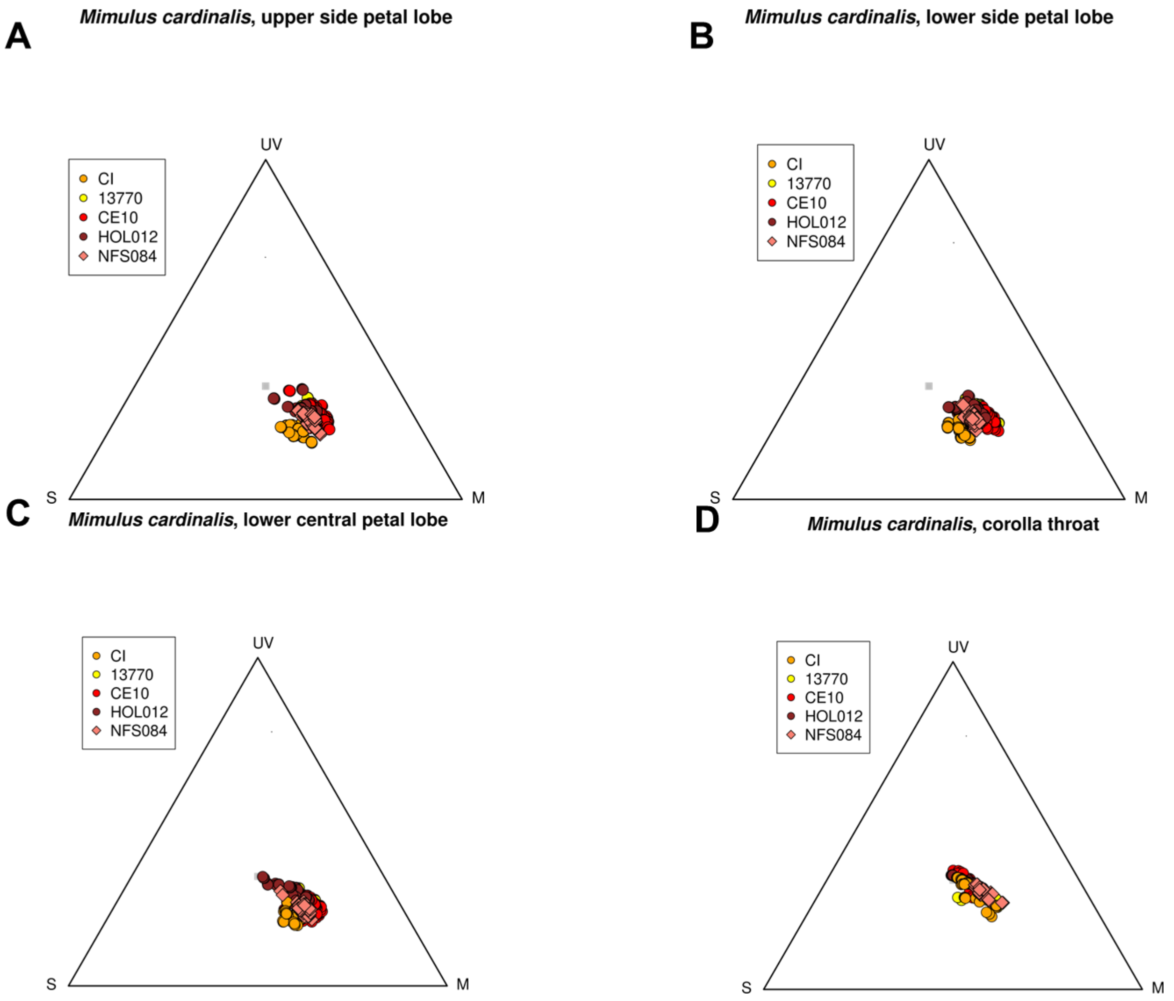


**Figure S4 -** Lepidopteran perception trichromat model based on spectral reflectance of (A) upper side petal lobes (B) lower side petal lobes (C) lower central petal lobes (D) corolla throat spots of M. cardinalis lines.

| Genome | Reference | % Mapped | % Properly Paired | % Singletons |
|---|---|---|---|---|
| 13770/SM | CE10 | 99.59 | 96.28 | 0.2 |
| CI | CE10 | 96.99 | 89.9 | 0.38 |

**Table S1** – Alignment scores of whole genomes of CI and 13770 when aligned with CE10.

| **Compound name** | **Kovats Retention Index (KRI)** | **NFS084 emission (ng/flower/hour ± SEM) (n = 15)** | **13770 emission (ng/flower/hour ± SEM) (n = 15)** | **CE10 emission (ng/flower/hour ± SEM) (n = 24)** | **HOL012 emission (ng/flower/hour ± SEM) (n = 15)** | **CI emission (ng/flower/hour ± SEM) (n = 15)** |
|---|---|---|---|---|---|---|
| **TOTAL VOLATILE EMISSIONS (58 COMPOUNDS)** | - | 38.787 ± 7.564 | 8.419 ± 0.812 | 3.314 ± 0.313 | 246.155 ± 24.579 | 28.997 ± 2.840 |
| **AROMATICS (4 COMPOUNDS)** | **-** | **-** | **-** | **1.251 ± 0.111** | **5.892 ± 0.674** | **0.832 ± 0.177** |
| Styrene[B] | 889 | - | - | - | 1.779 ± 0.324 (11) | - |
| Cymene[A] | 1021 | - | - | - | 2.137 ± 0.295 (15) | - |
| Phenylethyl alcohol[A] | 1113 | - | - | - | 1.201 ± 0.155 (14) | - |
| Methyl salicylate[A] | 1190 | - | - | 1.251 ± 0.111 (22) | 0.775 ± 0.117 (12) | 0.832 ± 0.177 (10) |
| **TERPENOIDS (27 COMPOUNDS)** | - | **32.361 ± 7.027** | **1.120 ± 0.236** | **1.123 ± 0.184** | **230.667 ± 23.481** | **9.682 ± 1.095** |
| α-Thujene[A] | 924 | - | - | - | 13.713 ± 2.065 (15) | - |
| α-Pinene[A] | 930 | 23.633 ± 5.221 (11) | - | - | 101.871 ± 10.234 (15) | - |
| Camphene[A] | 942 | - | - | - | 0.509 ± 0.096 (13) | - |
| Sabinene[A] | 970 | 4.814 ± 1.003 (15) | - | - | 53.515 ± 6.482 (15) | - |
| β-Pinene[A] | 971 | 3.142 ± 0.673 (13) | - | - | 10.215 ± 0.646 (15) | - |

| | | | | | | |
|---|---|---|---|---|---|---|
| 6-Methyl-5-hepten-2-one[A] | 987 | - | - | - | - | 8.096 ± 0.993 (15) |
| β-Myrcene[A] | 990 | - | - | - | 0.462 ± 0.070 (13) | - |
| β-Phellandrene[A] | 1024 | - | - | - | 13.775 ± 1.845 (15) | - |
| Eucalyptol[A] | 1027 | - | - | - | 0.718 ± 0.084 (15) | - |
| γ-Terpinene[A] | 1055 | - | - | - | 0.261 ± 0.070 (12) | - |
| Sabinene hydrate[A] | 1064 | - | - | - | 5.768 ± 0.693 (14) | - |
| (*Z*)-Linalool oxide[A] | 1070 | - | 0.227 ± 0.049 (10) | 0.663 ± 0.091 (20) | - | 0.929 ± 0.139 (14) |
| (*E*)-Linalool oxide[A] | 1086 | - | - | 0.460 ± 0.100 (17) | - | 0.657 ± 0.106 (14) |
| Pinene oxide[A] | 1092 | - | - | - | 3.505 ± 0.371 (15) | - |
| Unknown monoterpenoid[C] | 1096 | 0.183 ± 0.041 (10) | - | - | 0.414 ± 0.050 (14) | - |
| α-Campholenal[B] | 1122 | - | - | - | 0.324 ± 0.035 (15) | - |
| Unknown monoterpenoid[C] | 1129 | - | - | - | 0.198 ± 0.035 (12) | - |
| Nopinone[A] | 1132 | - | - | - | 3.131 ± 0.453 (15) | - |
| (*E*)-Verbenol[A] | 1143 | 0.590 ± 0.155 (10) | - | - | - | - |
| Sabine ketone[B] | 1151 | - | - | - | 10.902 ± 1.594 (15) | - |
| Pinocarvone OR Sabinone[B] | 1156 | - | - | - | 0.503 ± 0.111 (10) | - |
| Cryptone[B] | 1182 | - | - | - | 0.818 ± 0.140 (14) | - |

| | | | | | | |
|---|---|---|---|---|---|---|
| Myrtenol[A] | 1192 | - | - | - | 0.328 ± 0.050 (12) | - |
| Copaene[B] | 1368 | - | - | - | 2.037 ± 0.348 (15) | - |
| Dihydroionone (α or γ)[B] | 1391 | - | - | - | 5.612 ± 0.767 (15) | - |
| Valencene[A] | 1480 | - | 0.893 ± 0.206 (12) | - | 1.774 ± 0.444 (12) | - |
| Isoshyobunone[B] | 1543 | - | - | - | 0.315 ± 0.066 (11) | - |
| **FATTY ACID-DERIVED COMPOUNDS (9 COMPOUNDS)** | - | **3.080 ± 0.320** | **4.320 ± 0.329** | **0.941 ± 0.096** | **4.642 ± 0.482** | **3.365 ± 0.669** |
| 5-Hexenal, 4-methylene[B] | 897 | - | - | - | 0.563 ± 0.121 (12) | - |
| 1-Octen-3-one[B] | 977 | - | - | - | - | 1.102 ± 0.248 (12) |
| 1-Octen-3-ol[A] | 980 | - | - | - | 1.407 ± 0.180 (15) | 2.263 ± 0.429 (14) |
| 3-Octanone[B] | 987 | - | - | - | 1.082 ± 0.153 (13) | - |
| 3-Octanol[B] | 997 | - | - | - | 1.498 ± 0.234 (14) | - |
| Butanoic acid, 2 or 3-methyl, 3-hexenyl ester | 1232 | 0.450 ± 0.109 (10) | - | - | - | - |
| Hexyl 2-methylbutyrate[A] | 1238 | 2.326 ± 0.206 (15) | 4.072 ± 0.285 (15) | 0.661 ± 0.085 (21) | - | - |
| Phenethyl 2-methylbutyrate[A] | 1484 | 0.304 ± 0.059 (12) | 0.248 ± 0.060 (10) | 0.280 ± 0.033 (21) | - | - |
| Unknown alkane[C] | 1645 | - | - | - | 0.093 ± 0.016 (12) | - |

| | | | | | | |
|---|---|---|---|---|---|---|
| **OTHER COMPOUNDS (5 COMPOUNDS)** | - | **1.498 ± 0.203** | **1.999 ± 0.246** | **-** | **1.134 ± 0.220** | **3.418 ± 0.489** |
| Cyclopentane, *N,N*-dimethyl[B] | 694 | - | - | - | 0.867 ± 0.195 (10) | - |
| 3-Methylcyclopentyl acetate[B] | 901 | - | - | - | - | 0.852 ± 0.071 (15) |
| Lavender lactone[B] | 1041 | 1.193 ± 0.180 (15) | 1.801 ± 0.228 (15) | - | - | 2.566 ± 0.445 (15) |
| (*Z*)-Arbusculone[B] | 1052 | 0.305 ± 0.042 (13) | 0.198 ± 0.044 (10) | - | - | - |
| 1-Phenyl-1-butene OR Indan, 1-methyl[B] | 1080 | - | - | - | 0.268 ± 0.053 (11) | - |
| **UNKNOWN COMPOUNDS (13 COMPOUNDS)** | - | **1.848 ± 0.441** | **0.979 ± 0.099** | **-** | **3.819 ± 0.388** | **11.701 ± 1.096** |
| Unknown 1[C] | 728 | - | - | - | - | 0.729 ± 0.181 (10) |
| Unknown 2[C] | 929 | - | - | - | - | 6.437 ± 0.572 (15) |
| Unknown 3[C] | 949 | 0.486 ± 0.096 (10) | 0.715 ± 0.093 (13) | - | - | 3.115 ± 0.348 (14) |
| Unknown 4[C] | 968 | - | - | - | - | 0.251 ± 0.042 (12) |
| Unknown 5[C] | 1148 | - | - | - | - | 0.351 ± 0.064 (12) |
| Unknown 6[C] | 1226 | - | - | - | - | 0.686 ± 0.172 (12) |
| Unknown 7[C] | 1272 | 1.183 ± 0.393 (11) | - | - | 1.124 ± 0.196 (15) | - |
| Unknown 8[C] | 1312 | - | - | - | 0.269 ± 0.044 (12) | - |

| | | | | | | |
|---|---|---|---|---|---|---|
| Unknown 9[C] | 1393 | - | - | - | 0.743 ± 0.096 (15) | - |
| Unknown 10[C] | 1466 | - | - | - | 1.200 ± 0.188 (15) | - |
| Unknown 11[C] | 1685 | 0.179 ± 0.042 (10) | 0.264 ± 0.026 (14) | - | 0.483 ± 0.136 (11) | - |
| Unknown 12[C] | 1873 | - | - | - | - | 0.131 ± 0.026 (10) |

**Table S2 -** Volatile emissions of five lines of Mimulus cardinalis. SEM: standard error of the mean. Dash: compound was absent in this line. Numbers in parentheses after each emission value for individual compounds indicate the number of samples a compound was found in from that line (total sample numbers for each line are in the table header). Under the "Compound name" header, superscript letters: A: compound identity validated using authentic reference standards; B: compound identity validated using published Kovats Retention Indices, our calculated Kovats Retention Indices, and NIST Library spectrum matching; C: compound identity could not be validated and compound is listed as an unknown. Note for Cymene: m-/o-/p- structure could not be differentiated with authentic reference standards.
